# Anionic Lipid Trafficking and Cell Mechanics Regulate Membrane Electrical Potential in Non-Excitable Tissue Cells

**DOI:** 10.1101/2025.11.25.690536

**Authors:** Zhuoxu Ge, Qin Ni, Yuqing Yan, Yufei Wu, Bishwa Ranjan Si, Jinyu Fu, Dingchang Lin, Konstantinos Konstantopoulos, Yizeng Li, Sean X. Sun

## Abstract

The membrane potential of eukaryotic cells, classically described by the Goldman-Hodgkin-Katz model, is conventionally attributed to the steady-state balance of transmembrane ionic fluxes. Here, we demonstrate that non-excitable cells maintain stable, spatially heterogeneous membrane voltage gradients. Using the ratiometric voltage indicator JEDI-2P-cyOFP1, we quantitatively map membrane potential in live cells and find that cellular protrusions are consistently depolarized, whereas lateral membrane regions are hyperpolarized. These voltage gradients strongly correlate with anisotropic distributions of anionic lipids, including phosphatidylserine (PS) and phosphatidylinositol 3,4,5-trisphosphate (PIP_3_). Perturbation of cytoskeletal integrity, the ion exchanger NHE1, or vesicular trafficking disrupts both voltage anisotropy and lipid polarization, revealing an uncharacterized electromechanical regulatory mechanism. Moreover, membrane potential dynamically responds to cell shape, substrate stiffness, external electric fields, and cell cycle progression. To account for these phenomena, we develop a quantitative model that integrates ionic fluxes and lipid charge distributions, thereby explaining the emergence of voltage gradients. This framework establishes that membrane potential is governed not solely by ionic currents but also by lipid dynamics and mechanical signaling. Collectively, these findings identify membrane voltage as a global integrator of electrochemical and mechanical cues in eukaryotic cells.

## 2 Introduction

A defining property of eukaryotic cells is the transmembrane electrical potential [1, 2]. Charge separation across the plasma membrane arises from the capacitive nature of the lipid bilayer and the activity of ion channels and transporters sensitive to this potential. The interplay between membrane voltage and ion fluxes determines the cell’s electrical state. Rapid voltage dynamics (milliseconds to seconds) underlie action potentials and neuronal signaling, whereas slower changes (minutes to hours) influence cell migration [3, 4, 5], cell cycle progression [6, 7, 8, 9, 10, 2], and morphogenesis [11, 12]. Altered membrane potentials are also associated with disease states such as cancer [7, 13].

The Goldman-Hodgkin-Katz (GHK) model [14, 15] remains the prevailing quantitative framework for describing membrane potential, attributing voltage generation solely to ionic fluxes. This model assumes rapid intracellular ion diffusion, predicting a spatially uniform potential across the cell. Here, we show that this view is incomplete. Using a ratiometric live-cell voltage reporter, we reveal sustained membrane voltage gradients along the cell periphery. These spatial voltage patterns arise from the coordinated action of the cytoskeleton, ion transporters, and spatially heterogeneous lipid trafficking. Understanding how these systems collectively regulate potential distribution in non-excitable cells is essential for a comprehensive biophysical model of membrane voltage regulation.

To investigate this, we employ the genetically encoded voltage sensor JEDI-2P [16], using a ratiometric variant that corrects for variations in membrane and protein content along the cell boundary. We find that both inhibition of ion channels and disruption of the cytoskeleton significantly alter membrane potential. Moreover, mechanical perturbations – including cell spreading on adhesive substrates and externally applied stretch – induce pronounced voltage changes, indicating strong electromechanical coupling in live cells. The observed voltage gradients coincide with spatial variations in anionic lipid distribution. In particular, anisotropic trafficking of phosphatidylserine (PS) [17, 18] and phosphatidylinositol (PI) [18] establishes and maintains these gradients. Ion exchangers such as NHE1, together with the F-actin cytoskeleton, mediate lipid trafficking and contribute to the polarized voltage distribution. External electric fields further modulate membrane voltage patterns and cell polarization, while disruption of lipid trafficking abolishes galvanotaxis.

Over longer timescales, the membrane potential varies through the cell cycle, with gradual hyperpolarization during interphase and rapid depolarization during mitosis. Cell cycle arrest – whether by growth factor withdrawal or senescence – produces depolarization, suggesting that membrane voltage is a marker and regulator of proliferative state. Collectively, our results demonstrate that membrane potential is sustained by a dynamic interplay among ion transport, lipid trafficking, and cytoskeletal organization. A quantitative model incorporating ion fluxes and anionic lipid dynamics reproduces the observed voltage gradients, providing a unified framework for understanding the electromechanical regulation of membrane potential in eukaryotic cells.

## 3 Results

### 3.1 Measuring membrane voltage using ratiometric indicator JEDI-2P-cyOFP1

To investigate how membrane potential is regulated in non-excitable cells, we employed a ratiometric variant of the genetically encoded voltage indicator JEDI-2P [16]. JEDI-2P detects voltage changes through a voltage-sensing domain (VSD) fused to a circularly permuted green fluorescent protein (cpGFP). During depolarization, the VSD moves outward, reducing cpGFP fluorescence, whereas hyperpolarization drives inward motion and increases fluorescence. To correct for imaging artifacts and membrane morphology effects, JEDI-2P-cyOFP1 includes a reference fluorophore, cyOFP1, whose emission is voltage-insensitive. JEDI-2P-cyOFP1 was expressed in NIH 3T3 and HT1080 fibrosarcoma cells under a doxycycline-inducible promoter (Materials and Methods). The cpGFP/cyOFP1 fluorescence ratio thus provides a reliable readout of membrane potential dynamics in live cells (Fig. 1A,B).

**Figure 1:**
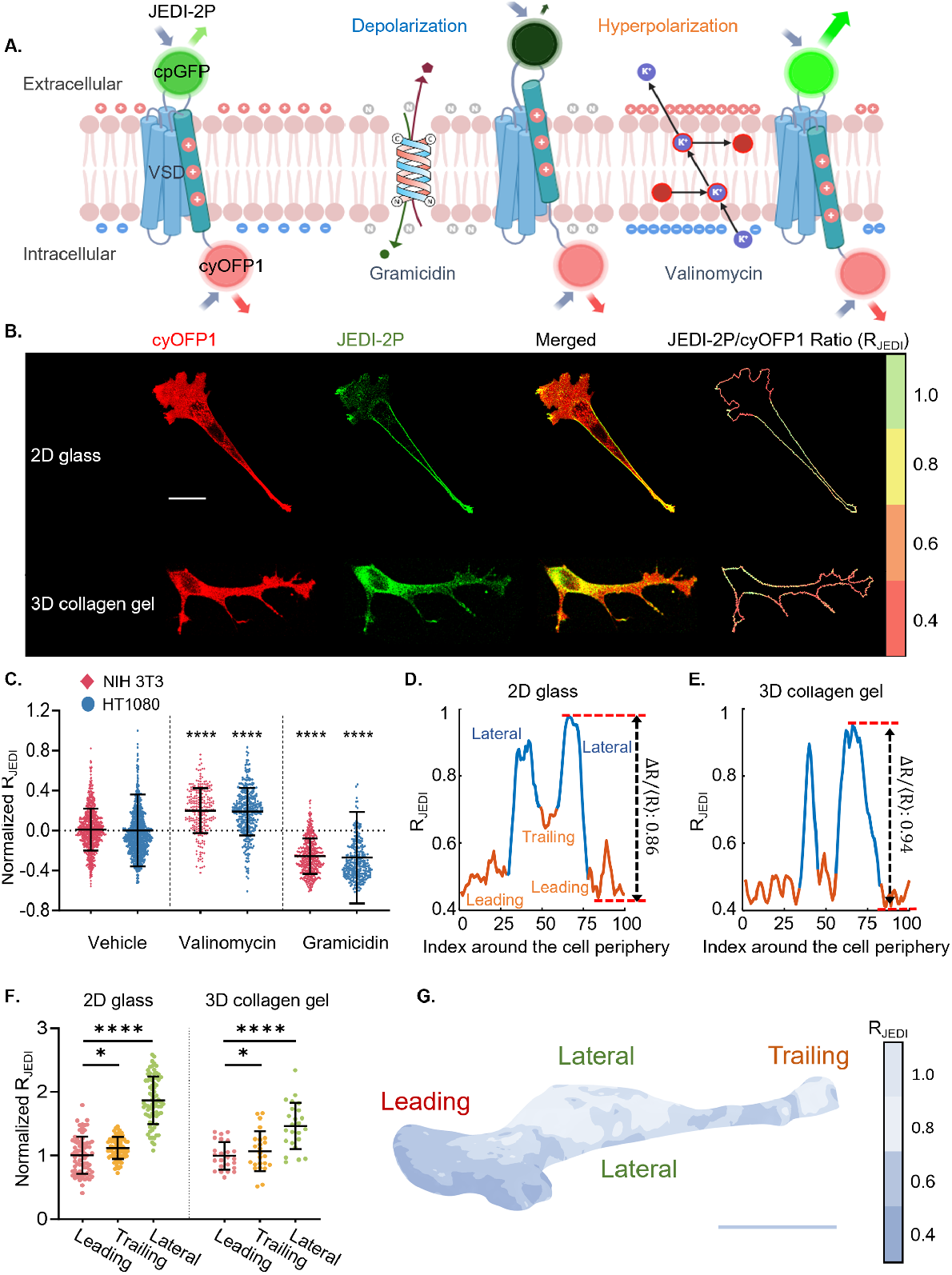
Spatial distribution of membrane potential in non-excitable cells. **(A)** Schematic of JEDI-2P-cyOFP1, a genetically encoded voltage indicator comprising a voltage-sensing domain (VSD) fused to a circularly permuted GFP (cpGFP). A red fluorescent protein (cyOFP1) is co-expressed as a ratiometric reference. Membrane depolarization decreases cpGFP fluorescence (lower cpGFP/cyOFP1 ratio), whereas hyperpolarization increases it. Gramicidin induces depolarization by enhancing ion permeability, while valinomycin causes hyperpolarization via potassium efflux. **(B)** Representative confocal images of 3T3 cells expressing JEDI-2P-cyOFP1, with ratiometric voltage (R_JEDI_) mapped along the cell boundary (two pixels inside the cell edge). Top: 3T3 cell migrating on a 2D glass substrate. Bottom: a 3T3 cell migrating in a 2 mg/mL 3D collagen gel. **(C)** Quantification of normalized R_JEDI_ in 3T3 and HT1080 cells treated with vehicle (0.2% DMSO), valinomycin (Val, 10 *μ*M), and gramicidin (20 nM). *n*_3T3_ = 932, 533, 248, and *n*_HT1080_ = 944, 285, 350. **(D**,**E)** R_JEDI_ intensity profiles along the cell boundary from (B), Δ*R* = *R*_Max_ *− R*_Min_; *(R)* = average ratio over the entire boundary. (D) 2D glass substrate, (E) 2 mg/mL 3D collagen gel. **(F)** Normalized R_JEDI_ in leading, trailing, and lateral regions of 3T3 cells on 2D glass substrate and in 2 mg/mL 3D collagen gel. Regional selection and averaging methods are detailed in Fig. S3. *n*_2D_ = 76, *n*_3D_ = 24. **(G)** 3D reconstruction of membrane voltage in a 3T3 cell. Deconvoluted images for each optical section are shown in Fig. S2C. **(C, F)** Error bars represent standard deviation. (C) Kruskal-Wallis tests followed by Dunn’s post hoc multiple-comparisons test vs. vehicle controls. (F) One-way ANOVA with Dunnett’s test vs. leading-edge region. **(B, G**) Scale bar: 20 *μ*m.

To quantify voltage changes, we segmented the cell boundary and computed the cpGFP/cyOFP1 ratio along the entire perimeter (Fig. 1B), yielding both spatial voltage profiles and average potentials per cell. We validated the sensor using the ionophores gramicidin and valinomycin. Gramicidin forms nonselective ion channels that dissipate negative charge and depolarize the membrane [19, 10], producing a decrease in the fluorescence ratio (Fig. 1A,C). Conversely, valinomycin increases potassium permeability [20, 21], hyperpolarizing the membrane and increasing the ratio (Fig. 1A,C). These results confirm that JEDI-2P-cyOFP1 quantitatively reports membrane potential changes in non-excitable cells.

We further assessed the stability of the cpGFP/cyOFP1 ratio under varying conditions (Fig. S1A-F). The ratio was independent of expression level, as shown by the lack of correlation between cyOFP1 intensity and the cpGFP/cyOFP1 ratio across cells (Fig. S1A,D). After doxycycline removal, cyOFP1 intensity gradually declined due to protein turnover (Fig. S1B,E), yet the fluorescence ratio remained constant over 12 hours (Fig. S1C,F). Together, these results demonstrate that JEDI-2P-cyOFP1 provides a robust and quantitative measure of membrane potential, stable across expression levels and over extended imaging periods.

### 3.2 Spatially distributed membrane voltage gradient

We next examined the spatial organization of membrane potential in live cells. Imaging with JEDI-2P-cyOFP1 revealed a strikingly non-uniform distribution of membrane voltage (reported as the cpGFP/cyOFP1 ratio) along the cell periphery in both 3T3 and HT1080 cells. A similar voltage gradient is observed in a recent study in neutrophils using a different ratiometric voltage probe [22]. In 3T3 cells, the leading edge was markedly depolarized, whereas the lateral regions were more hyperpolarized (Fig. 1D,F). This gradient was not due to uneven membrane distribution of JEDI-2P (Fig. S1J). Although cyOFP1 fluorescence appeared enriched at the trailing edge, this reflected local cell contraction and did not correspond to an increased cpGFP/cyOFP1 ratio. To quantify spatial variation, we defined a “voltage uniformity” metric: uniformity = (*R*_Max_ −*R*_Min_)*/* ⟨*R* ⟩, where larger values indicate greater voltage heterogeneity.

HT1080 cells displayed similar voltage non-uniformity, though with smaller differences between protrusive and lateral regions (Fig. S1I), despite comparable mean membrane potentials (Fig. S1H). Interestingly, both leading and trailing edges were depolarized, but the trailing edge was slightly more hyperpolarized than the leading edge. Because ions diffuse rapidly in the cytoplasm, electrical potential should be uniform throughout the cell interior [23, 24, 25]. The observed gradients therefore likely reflect JEDI’s sensitivity to the local transmembrane potential – i.e., the voltage difference across the lipid bilayer – rather than variations in the bulk cytoplasmic potential (Fig. 2A). This local potential is fundamentally more important since transmembrane proteins are sensitive to the local membrane potential, as detected by JEDI.

**Figure 2:**
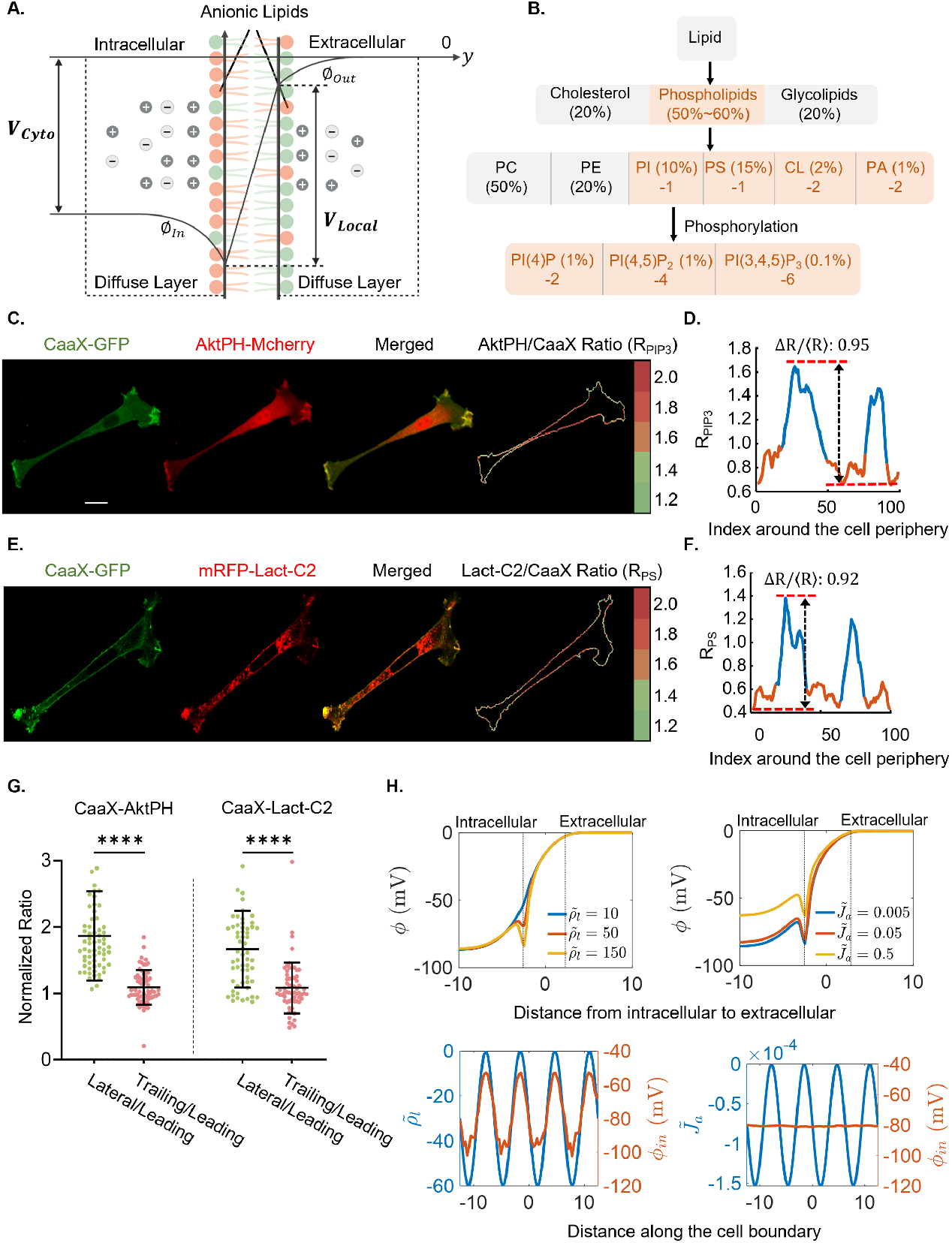
Membrane potential distribution is governed by anionic lipid asymmetry. **(A)** Schematic illustrating membrane potential components. The local membrane potential (*V*_Local_) represents the voltage drop from the outer (*φ*_Out_) to inner leaflet (*φ*_In_) membrane leaflet, while the cytoplasmic potential (*V*_Cyto_) denotes the voltage difference between intraand extracellular fluids. **(B)** Summary of plasma membrane lipid composition and corresponding charge contributions[30, 32, 33, 34, 35, 31, 36, 37, 38, 39, 40]. **(C, D)** Representative confocal imaging (C) of 3T3 cells co-expressing AktPHmCherry (PIP_3_ sensor) and CaaX-GFP (membrane marker), with the calculated AktPH/CaaX Ratio (*R*_PIP3_) mapped along the cell boundary. (D) *R*_PIP3_ intensity profile along the cell boundary. **(E, F)** Representative confocal imaging (E) of 3T3 cells co-expressing mRFP-Lact-C2 (PS sensor) and CaaX-GFP, with the calculated Lact-C2/CaaX Ratio (*R*_PS_). (F) *R*_PS_ intensity profile along the cell boundary. **(G)** Quantification of normalized *R*_PIP3_ and *R*_PS_ across leading, lateral, and trailing regions in 3T3 cells. Regional selection and averaging are described in Fig. S2A. *n*_PIP3_ = 66, *n*_PS_ = 61. **(H)** Theoretical model predicts membrane potential distribution as a function of ion flux 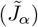 and anionic lipid concentration 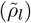. Both factors modulate local potential, but inner leaflet voltage variations primarily arise from anionic lipids. **(G)** Error bars represent standard deviation; Mann-Whitney tests were used for pair-wise comparisons. **(C, E)** Scale bar: 20 *μ*m.

To exclude geometric or optical artifacts, we also collected confocal z-stacks and applied RichardsonLucy deconvolution [26] to correct for wavelength-dependent microscope point spread function (Fig. S2, Materials and Methods). This reconstruction yielded high-fidelity 3D maps of membrane voltage (Fig. 1G). Cells embedded in 3D collagen gels (2 mg/mL) exhibited comparable spatial voltage gradients to those observed on 2D substrates (Fig. 1E,F).

### 3.3 Anionic lipid distribution governs local membrane potential

To understand the uneven JEDI-2P signals observed along the cell boundary, we hypothesized that this variation arises from a nonuniform charge distribution across the plasma membrane. The inner leaflet is enriched in negatively charged phospholipids [27], which attract counterions such as K^+^, while the outer leaflet contains highly charged glycoproteins [28, 29]. The overall electrical potential results from these fixed charges and the surrounding mobile ions. The potential profile *V* (*y*) across the membrane can be described by the Poisson equation (Supplementary Information, SI), where *y* denotes the coordinate normal to the membrane surface (Fig. 2A). In the extracellular region, *V* (*y*) approaches the bulk potential (~ 0 mV), whereas in the cytoplasm it reaches approximately −70 mV. Mobile ions form a Debye layer that interacts with lipid headgroup charges, shaping the local potential. Because negatively charged lipids are concentrated in the inner leaflet [27], they generate an additional potential drop (*φ*_In_). We define the cytoplasmic voltage *V*_Cyto_ as the bulk potential difference across the membrane, and the local membrane potential *V*_Local_ as the voltage difference between the inner and outer leaflets (SI). As a transmembrane reporter, JEDI-2P-cyOFP1 primarily senses *V*_Local_ (Fig. 2A).

To test this hypothesis, we examined which phospholipids contribute to membrane surface charge. The dominant species - phosphatidylcholine (PC, 40-50%) and phosphatidylethanolamine (PE, 15-25%) - are zwitterionic and electrically neutral [30, 31, 32, 33]. In contrast, minor anionic phospholipids such as cardiolipin (CL) and phosphatidic acid (PA) carry negative charges, though both are present in low abundance (≤ 1%) [34, 35]. The main contributors to the membrane’s negative charge are phosphatidylserine (PS, 20%) and phosphatidylinositol (PI, 12%) [31, 36]. PI can be phosphorylated into phosphatidylinositol 4-phosphate (PIP), phosphatidylinositol 4,5-bisphosphate (PIP_2_), and phosphatidylinositol 3,4,5-trisphosphate (PIP_3_), increasing its charge to −2, −4 and −6, respectively [37, 38, 39, 40]. Although these species are low in abundance (PIP ~1%, PIP_2_ ~1%, PIP_3_ ~0.1%), their high charge density makes them potent regulators of local voltage. We therefore focused on mapping PS and PIP_3_ distributions using live-cell biosensors (Fig. 2B).

PIP_3_ localization was visualized using the AktPH-mCherry probe, containing the pleckstrin homology domain of Akt1 fused to mCherry [41], co-expressed with CaaX-GFP to mark the membrane. The mCherry/GFP ratio along the boundary quantified PIP_3_ enrichment (Fig. 2C). For PS, we used a Lactadherin C2 (LactC2)-mCherry sensor [42] co-transfected with CaaX-GFP and analyzed the mCherry/GFP ratio using the same boundary-masking approach (Fig. 2E). Both PIP_3_ and PS probes displayed higher fluorescence ratios in the lateral membrane compared with protrusions (Fig. 2D,F,G), consistent with JEDI-2P patterns. Note that due to membrane ruffling, raw AktPH-mCherry intensities can be high in the protrusion regions (Fig. S3I), however the ratio showed that PIP3 is lower. Moreover, whereas JEDI signals showed slightly greater hyperpolarization at the trailing edge (Fig. 1F, 2G), PIP_3_ and PS distributions did not show this asymmetry.

These results support the interpretation that JEDI-2P-cyOFP1 primarily reports the local membrane potential, which correlates with the spatial distribution of anionic lipids. Regions enriched in PS and PIP_3_ – where negative charge density is highest – correspond to more hyperpolarized membrane domains. To further explore this mechanism, we developed a two-dimensional Poisson-Nernst-Planck model that includes the membrane region and spans from the cytoplasm to the extracellular space (SI). Mobile ions (Na^+^, K^+^, Cl^*−*^, etc) diffuse and traverse ion channels under steady-state conditions. Simulations show that ionic fluxes alone cannot generate substantial voltage differences along the cell boundary; instead, spatially heterogeneous distributions of negatively charged lipids are sufficient to produce the observed potential gradients. Even though diffusible cations are attracted to anionic lipids at the inner leaflet, due to screening (characterized by a screening Debye length), the negative charges are not completely neutralized (SI). The resulting voltage profile (Fig. 2H) demonstrates that localized enrichment of anionic lipids generates distinct hyperpolarized and depolarized membrane regions. Fig. 1F is consistent with the OEM prediction that asymmetric/polarized distribution of ion transporters will generate a slight voltage difference between the leading and trailing edges [43]. Together, these findings confirm that JEDI-2P-cyOFP1 detects local transmembrane potentials and that nonuniform lipid charge distributions are the primary drivers of membrane voltage anisotropy.

### 3.4 Ion fluxes modulate membrane potential

We next investigated how ionic fluxes influence local membrane potential in non-excitable cells. This question was motivated by the osmotic engine model (OEM), which posits that asymmetric ion fluxes drive water movement and cell migration [44]. OEM predicts a mild electrical gradient from the leading to trailing edge of migrating cells (Fig. 1F) [45], yet this mechanism cannot explain the large potential differences observed between protrusions and lateral regions. We focused on the Na^+^/H^+^ exchanger (NHE1), which regulates cell volume and migration by promoting local swelling and is enriched at the leading edge [46, 47, 45].

Surprisingly, inhibition of NHE1 with EIPA caused strong cell hyperpolarization in both 3T3 and HT1080 cells (Fig. 3A, S4B), despite the electroneutral nature of sodium influx balanced by proton efflux [48]. Separate imaging of cyOFP1 after EIPA treatment confirmed that this hyperpolarization was not due to cyOFP1 pH sensitivity (Fig. S4E). The response was most pronounced at protrusions during the first 5 minutes of EIPA treatment, then gradually returned to baseline over 30 minutes (Fig. 3B, D). Immunofluorescence revealed higher NHE1 enrichment in protrusive regions (Fig. 3F) [49, 50], consistent with stronger local responses. Inhibiting NHE1 also reduced voltage gradients, equalizing potential across the cell boundary (Fig. S4A).

**Figure 3:**
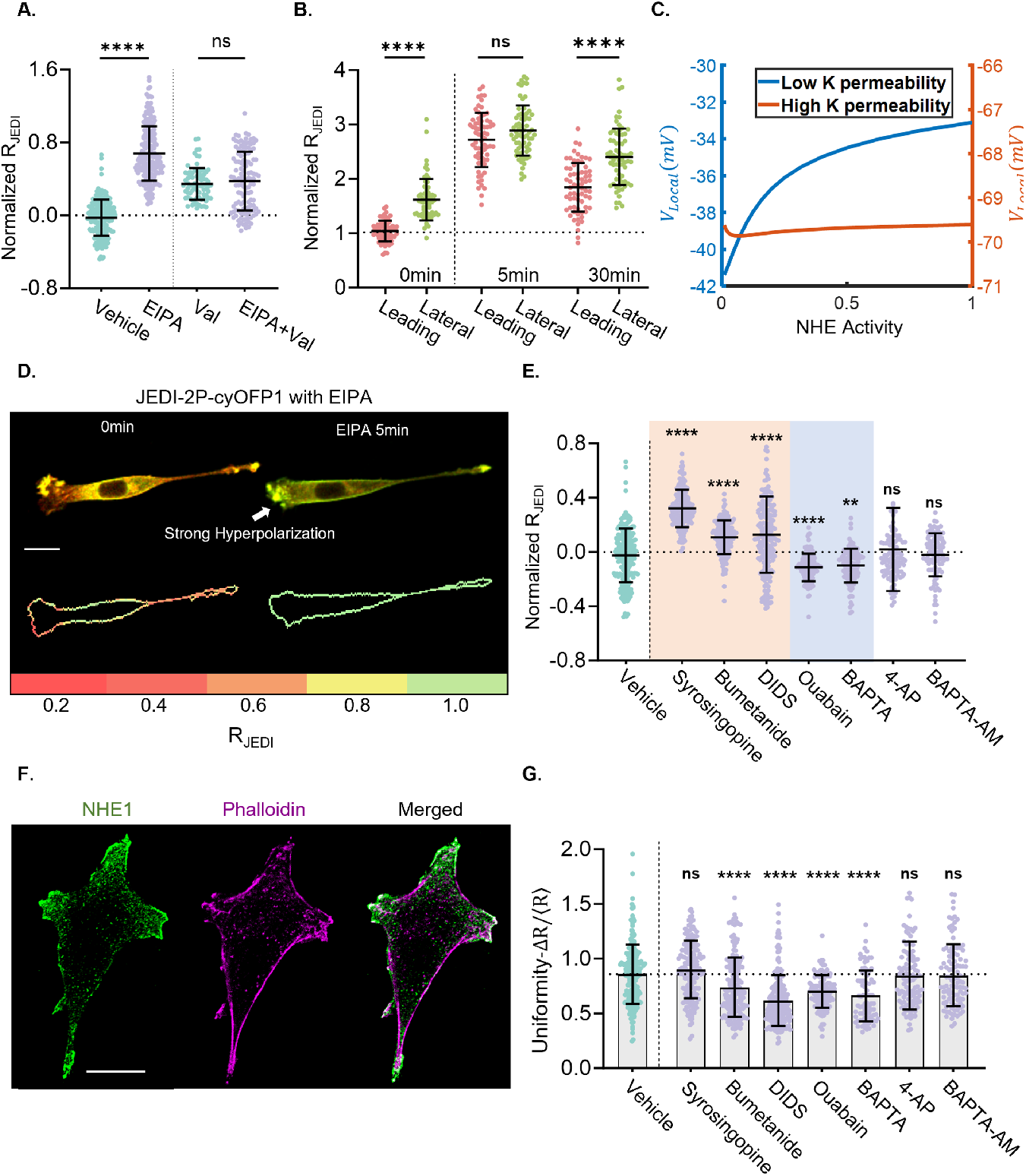
Influence of ionic fluxes on the membrane potential. **(A)** Normalized *R*_JEDI_ changes in 3T3 cells treated with vehicle control (0.2% DMSO), EIPA (40 *μ*M), valinomycin (Val), or Val + EIPA. *n*_3T3_ = 200, 200, 78, 122. **(B)** Time-dependent changes in *R*_JEDI_ in 3T3 cells, comparing the leading and lateral regions after NHE inhibition. *n*_3T3_ = 68. **(C)** Model prediction of local membrane voltage *V*_Local_ as a function of NHE activity. Orange: high K^+^ permeability; blue: low K^+^ permeability. **(D)** Representative confocal images of 3T3 cells expressing JEDI-2P-cyOFP1 before and after EIPA treatment (0 and 5 min), with mapped *R*_JEDI_ along the cell boundary. **(E)** Normalized *R*_JEDI_ changes in 3T3 cells treated with ion transport inhibitors: vehicle control, Syrosingopine (10 *μ*M; MCT4 transporter inhibitor), Bumetanide (40 *μ*M; NKCC inhibitor), DIDS (40 *μ*M; Cl^*−*^/HCO^*−*^_3_ exchanger inhibitor), BAPTA (1.2 mM, extracellular Ca^2+^ chelator), ouabain (100 *μ*M, NKA inhibitor), 4-aminopyridine (4-AP, 4 mM, voltage-gated K^+^ channel blocker), and BAPTA-AM (10 *μ*M, intracellular Ca^2+^ chelator). *n*_3T3_ = 200, 153, 165, 200, 120, 112, 110, 103. **(F)** Representative immunofluorescence images showing NHE1 (green) and F-actin (phalloidin; magenta) in 3T3 cells. **(G)** Quantification of membrane potential uniformity in 3T3 cells under ion transport treatments shown in (E). **(A, B, E, G)** Error bars represent standard deviation. (A, B) Mann-Whitney tests; (E, G) Kruskal-Wallis tests followed by Dunn’s multiple-comparisons test versus vehicle control. **(D, F)** Scale bar: 20 *μ*m.

To understand this effect mechanistically, we developed a mathematical model incorporating Na^+^, K^+^, Cl^*−*^, and H^+^ fluxes through major channels, co-transporters, and exchangers (SI). The model predicts that decreased NHE1 activity reduces intracellular Na^+^, thereby reducing Na^+^/K^+^-ATPase (NKA) flux and decreasing intracellular K^+^, leading to hyperpolarization. This prediction was validated experimentally: pretreatment with valinomycin, which increases K^+^ permeability, diminished EIPA-induced hyperpolarization (Fig. 3A, C). Thus, NHE1 indirectly modulates membrane potential via Na^+^ homeostasis, in addition to its role in osmotic-driven protrusion formation. Blocking ezrin, which links NHE1 to the actin cytoskeleton [47, 49], also induced hyperpolarization, though less than direct NHE1 inhibition (Fig. S4C), suggesting that cytoskeletal coupling enhances NHE1 function.

We next examined how other ionic perturbations influence membrane potential. Removing extracellular Ca^2+^ with BAPTA caused strong depolarization, whereas chelating intracellular Ca^2+^ with BAPTA-AM had no effect. Inhibition of the Na^+^-K^+^-2Cl^*−*^ cotransporter with bumetanide and blockade of 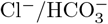 transporter with DIDS both caused hyperpolarization by reducing cation influx. Inhibition of lactate-proton transporter MCT with syrosingopine also caused hyperpolarization. Conversely, inhibition of NKA with ouabain led to depolarization by collapsing ionic gradients. Blocking voltage-gated K^+^ channels with 4-aminopyridine had little effect (Fig. 3E, S4B). The effects of bumetanide and ouabain closely matched model predictions (Fig. S4D), supporting the model’s validity.

Interestingly, several of these treatments also reduced spatial voltage heterogeneity. Inhibiting NHE1 (EIPA), NKA (ouabain), NKCC (bumetanide), anionic exchange (DIDS), or removing extracellular Ca^2+^ (BAPTA) all increased voltage uniformity (Fig. 3G). This suggests that specific ion transporters interact with anionic lipids – particularly PIP_3_ and PS – to shape local membrane potential. Previous studies emphasized lipid regulation of ion channels via PIP_2_ [51, 52]; our findings suggest a reciprocal mechanism in which ion fluxes also influence anionic lipid distribution, forming a feedback loop that maintains membrane voltage gradients.

Finally, we examined the effects of PI3K inhibition on membrane potential. In HT1080 cells, the PI3K inhibitor LY294002 (LY29) caused mild depolarization, consistent with reduced negative surface charge (Fig. S4G). In contrast, 3T3 cells exhibited hyperpolarization, which was abolished by NHE1 inhibition (Fig. S4F). LY29 also acidified 3T3 cells but not HT1080 cells, indicating that suppressed NHE1 activity underlies the hyperpolarization. This cell-type-specific response reflects differential coupling of the PI3K-Akt-NHE1 pathway: in 3T3 cells, PI3K/Akt inhibition reduces NHE1 activity and cytoplasmic pH [53, 54], whereas HT1080 cells lack this compensatory regulation.

### 3.5 Membrane trafficking regulates anionic lipid distribution and membrane potential gradient

To investigate the origin of the uneven anionic lipid distribution, we hypothesized that regional differences in membrane trafficking and endocytosis contribute to the observed lipid asymmetry. Using Fluorescence Recovery After Photobleaching (FRAP) with the PS reporter, we selectively bleached lateral and protrusive regions and monitored fluorescence recovery. The lateral membrane exhibited a higher recovery plateau than protrusions (Fig. 4A), indicating enhanced membrane turnover. A mathematical model incorporating lipid diffusion and trafficking (SI) reproduced these dynamics, suggesting higher net rates of vesicle import, and fusion/endocytosis activity in the lateral regions.

**Figure 4:**
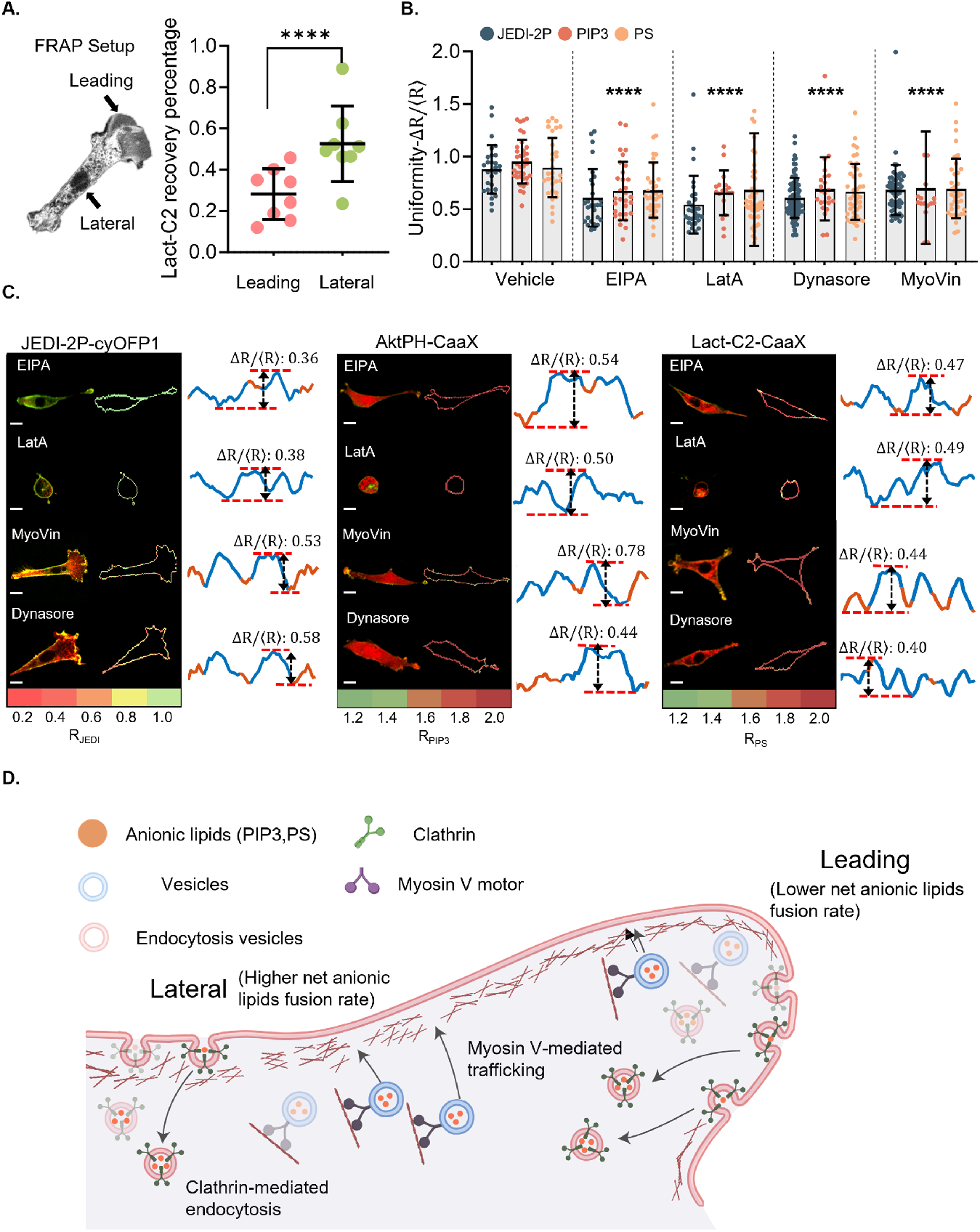
Membrane trafficking regulates anionic lipid distribution and membrane potential. **(A)** Fluorescence Recovery After Photobleaching (FRAP) of PS mobility in 3T3 cells. Left: Experimental setup showing bleached regions at the leading and lateral regions. Right: Quantification of LactC2 recovery kinetics. *n*_3T3_ = 8. **(B)** Quantification of voltage, PIP_3_, and PS uniformity in 3T3 cells treated with trafficking inhibitors: vehicle control, EIPA, LatA (2 *μ*M), Dynasore (80 *μ*M; dynamin inhibitor), and MyoVin (80 *μ*M; myosin-V inhibitor). *n*_JEDI_ = 29, 28, 29, 87, 55. *n*_PIP3_ = 37, 31, 18, 23, 16. *n*_PS_ = 31, 36, 47, 42, 37. **(C)** Representative confocal images of 3T3 cells expressing JEDI-2P-cyOFP1, AktPH-CaaX, and Lact-C2-CaaX, with calculated ratio maps (*R*) along the cell boundary. Corresponding distributions under inhibitor treatments are shown in (B). **(D)** Schematic illustrating vesicle trafficking differences between the lateral and protrusion regions. The lateral region exhibits higher trafficking activity than the leading edge. **(A, B)** Error bars represent standard deviation. (A) Mann-Whitney tests were used for two-condition comparisons. (B) Kruskal-Wallis tests with Dunn’s multiple-comparisons test were applied against the vehicle control. Statistical significance was consistent across indicators (JEDI-2P, PIP_3_, and PS) for each treatment. **(C)** Scale bar: 20 *μ*m.

We next examined how disrupting cell polarity and vesicle trafficking affects both membrane potential and anionic lipid distribution. Inhibiting NHE1 with EIPA or depolymerizing F-actin with Latrunculin A (LatA) led to a more uniform distribution of voltage, PS, and PIP_3_ along the cell boundary, reflected by decreased JEDI-2P-cyOFP1 ratio variation (Fig. 4B,C). Similarly, blocking clathrin-mediated endocytosis with Dynasore [55] or myosin V-dependent vesicle transport with MyoVin-1 [56] produced comparable effects. These findings indicate that lateral regions undergo more active vesicle trafficking than protrusions, which contributes to the accumulation of anionic lipids and associated hyperpolarization in these zones (Fig. 4D).

To further assess the role of endocytosis, we performed FRAP following inhibition of trafficking pathways. EIPA pretreatment equalized recovery rates between lateral and protrusive regions, consistent with the loss of polarity and cell shrinkage. In contrast, Dynasore reduced recovery in both regions, with a stronger effect laterally (Fig. S5C,D), confirming that lateral membranes normally experience higher trafficking flux. Together, these results demonstrate that spatially heterogeneous membrane trafficking and endocytosis establish the asymmetric distribution of anionic lipids, thereby shaping voltage gradients along the cell periphery. If there is higher vesicle fusion at the cell lateral regions and higher endocytosis/membrane intake at the protrusion regions, this would generate accumulation/depletion of anionic lipids at the lateral/protrusion regions, respectively. This would explain the observed membrane voltage gradient.

### 3.6 Mechanical forces and cytoskeletal dynamics affect local membrane potential

Beyond ionic regulation, cytoskeletal dynamics exert a strong influence on membrane potential. In both 3T3 and HT1080 cells, membrane voltage changes were observed during cytoskeletal-driven events such as spreading and division. Upon detachment and reseeding, cells were initially hyperpolarized, then gradually depolarized as they spread and increased in surface area (Fig. S7A, D). A similar pattern occurred during mitosis: cells hyperpolarized during mitotic rounding and depolarized as daughter cells spread (Fig. S7B, E). Across these processes, membrane potential negatively correlated with cell area, indicating that spreading and surface expansion are associated with depolarization (Fig. S7C). Transient protrusions and retractions also produced rapid, reversible voltage shifts, highlighting tight coupling between cell mechanics and membrane voltage.

Mechanical forces and substrate stiffness further modulated cell potential [57]. When subjected to 20% axial stretch, 3T3 cells showed transient hyperpolarization followed by recovery, whereas HT1080 cells progressively depolarized (Fig. S7F), consistent with their distinct NHE1 activation patterns [47, 53]. Hypertonic shock produced similar effects: 3T3 cells briefly hyperpolarized, while HT1080 cells showed little response (Fig. S8A), indicating cell type-specific electromechanical coupling.

Substrate stiffness also strongly influenced steady-state membrane potential [58]. 3T3 cells cultured on softer substrates (15 kPa to 400 Pa) were more depolarized than those on stiffer surfaces such as glass (Fig. S7G). Long-term culture revealed a persistent “electromechanical memory”: cells cultured on 15kPa PDMS substrate and transferred to glass substrate remained depolarized, showing the same voltage as cells grown on 15kPa (Fig. S8B). Finally, actin depolymerization with Latrunculin A induced hyperpolarization and increased PIP_3_ levels in 3T3 cells, but had no effect in HT1080 cells (Fig. S8C,E).

Together, these results reveal a tight interplay between cytoskeletal organization, mechanical forces, ion exchangers, and anionic lipids in regulating membrane potential. This network is robust in 3T3 cells but appears rewired in HT1080 cells, underscoring distinct electromechanical coupling mechanisms between normal and transformed cell types.

### 3.7 External electrical field stimulation influences cell protrusion formation, membrane potential, and anionic lipid distribution

Over the course of the entire cell cycle, cells maintained a generally hyperpolarized state, with a notable voltage reset occurring at the M phase (Fig. S6D). To determine whether membrane potential dynamics reflect cell cycle progression, we arrested HT1080 cells in late G1 using the CDK4/6 inhibitor Palbociclib [59]. Although Palbociclib-treated cells continue to grow in size during G1 arrest [60], their membrane potential remained stable (Fig. S6E), indicating that voltage dynamics depend more on cell cycle state than on cell size. We next examined two non-proliferative states – quiescence and senescence – that represent distinct forms of cell cycle exit. Quiescence was induced by serum starvation [61], and senescence by Doxorubicin treatment [62]. Both conditions produced marked membrane depolarization relative to proliferating cells, consistent with prior reports of depolarization during cell cycle exit [63, 7, 64, 65] (Fig. S6A, C). Quiescent cells maintained normal morphology but exhibited reduced voltage anisotropy, displaying more uniform potentials across protrusive and lateral regions (Fig. S6B). Senescent cells, characterized by enlarged, flattened morphology with multiple protrusions, showed near-complete voltage uniformity, likely reflecting global depolarization. Together, these results demonstrate that membrane potential dynamics are tightly linked to cell cycle state and largely independent of cell size or shape.

Given that electric fields can guide migration in various cell types [66, 67, 43, 68, 69], we next investigated whether external fields modulate membrane voltage and lipid polarity. Using JEDI-expressing 3T3 cells, we mapped voltage distributions before and after applying a 3 V/cm direct current (DC) electric field (Fig. 5A). Prior to stimulation, cells exhibited random polarity. Upon field application, a depolarized region emerged at the field-facing side (0^*°*^), while a hyperpolarized region appeared at the rear (180^*°*^), indicating field-aligned voltage polarization.

**Figure 5:**
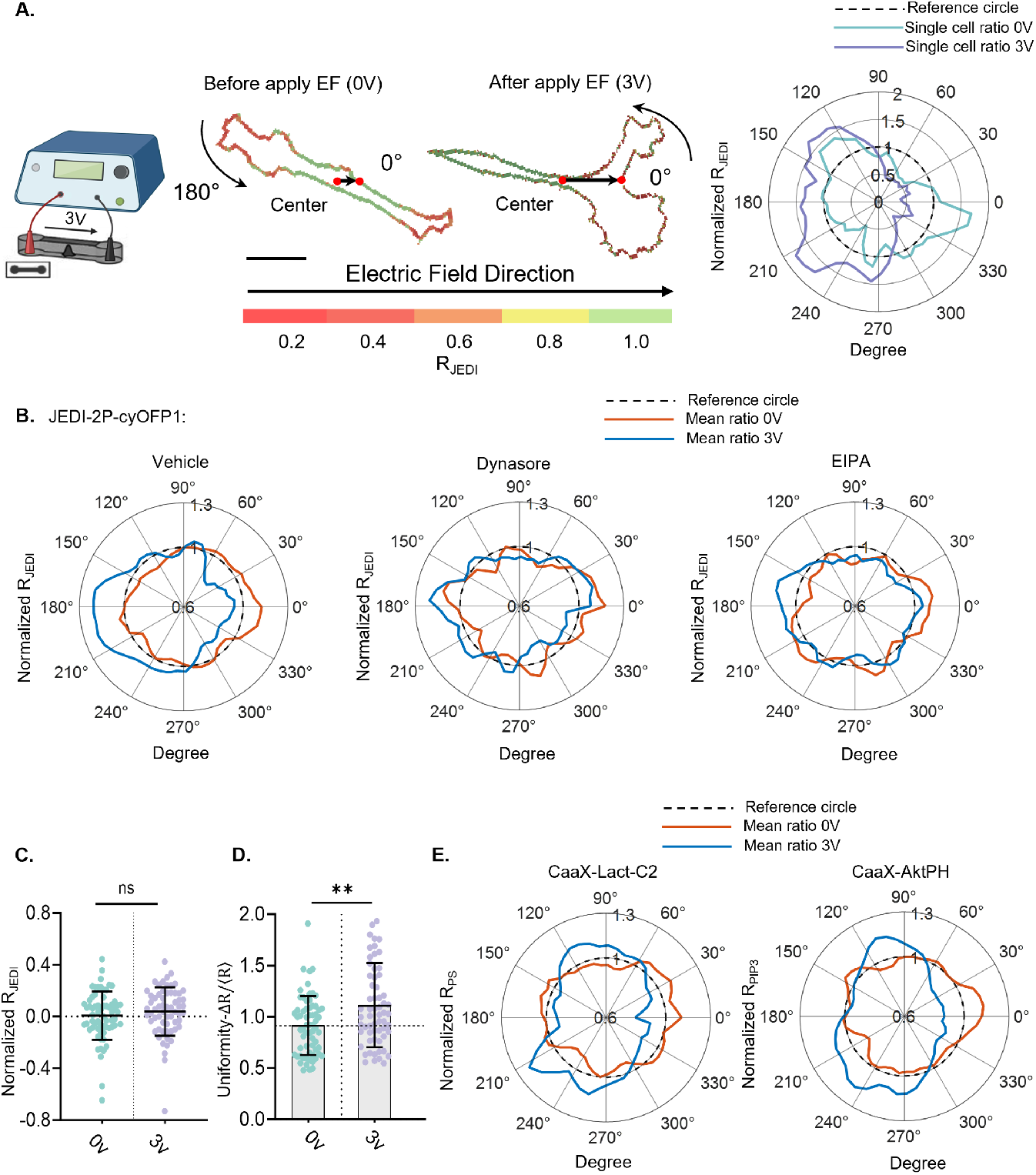
Membrane potential and anionic lipid redistribution in response to external electric fields. **(A)** Electrotaxis in 3T3 cells. Left: Schematic of the setup applying a 3 V/cm DC electric field (EF) in a microfluidic device. Middle: Representative *R*_JEDI_ images before and after EF application, showing cell polarity change. Cell boundaries are mapped in polar coordinates centered at the cell’s center of mass. 0^*°*^ is aligned with the EF direction. Right: Polar plots of single-cell *R*_JEDI_ distribution before and after EF application, normalized to each cell’s mean *R*_JEDI_. **(B)** Average spatial *R*_JEDI_ distribution in 3T3 cells treated with vehicle, Dynasore, or EIPA before and after EF application. *n*_3T3_ = 40, 36, 37. **(C)** Normalized *R*_JEDI_ changes in 3T3 cells before and after applying a 3 V/cm EF. *n*_3T3_ = 65. **(D)** Quantification of membrane potential uniformity in 3T3 cells before and after EF application. **(E)** Average spatial distribution of *R*_PS_ and *R*_PIP3_ in 3T3 cells before and after EF application. *n*_PS_ = 25, *n*_PIP3_ = 22. **(C, D)** Error bars represent standard deviation. Kruskal-Wallis tests with Dunn’s multiple-comparisons test were performed relative to the control. **(A)** Scale bar: 20 *μ*m

At the population level, normalized fluorescence ratios shifted from circular (isotropic) to asymmetric distributions, reflecting polarization toward the field (Fig. 5B). Although the mean membrane potential remained unchanged, voltage uniformity increased significantly following field exposure (Fig. 5C, D). Inhibiting NHE1 with EIPA or blocking endocytosis with Dynasore reduced field responsiveness from ~80% to ~40% (Fig. S6F) and attenuated polarity reorientation, implicating endocytic trafficking in electro-sensing.

Because voltage gradients coincide with polarized anionic lipid distributions, we examined PS and PIP3 reporters under electric field stimulation. Both lipids redistributed toward the field-facing side, mirroring the voltage polarity shift (Fig. 5E). Thus, external electric fields induce coordinated reorganization of membrane voltage and anionic lipids, a process requiring intact endocytic machinery.

## 4 Discussion and Conclusions

Together with the cell size, the cell electrical potential is a global property that potentially influences all aspects of cell function. In this paper, we showed that the membrane potential is determined by a combination of ion channel activity and anionic lipid trafficking. Trafficking of anionic lipids is anisotropic, leading to accumulation of lipids and membrane hyperpolarization at the cell lateral regions and depolarization at the cell leading edge. We showed that local membrane potential coincides with the asymmetric distribution of anionic lipids, particularly phosphatidylserine (PS) and phosphatidylinositols (PI). Since the cytoskeleton and ion exchangers, such as NHE1, are involved in membrane lipid trafficking, inhibiting the cytoskeleton, NHE1, and ezrin will influence membrane potential polarization. Therefore, there is extensive electromechanical coupling in the cell. Deforming the cell will influence cell electrical activity and vice versa. Since ion exchangers such as NHE1 also influences cell pH and metabolism, cell mechanical behavior will influence cell metabolic activity. Thus, a network of interactions, which also includes canonical signaling pathways such as MAPK/PI3K/Akt, is emerging that determines the overall state. Membrane potential, together with ion channels interacting with the actin cytoskeleton, likely provide global signals that influence the activity of these signaling pathways. This also implies that protein synthesis/growth and lipid metabolism are likely regulated together, which is consistent with the “scaling” idea, where all cell components, proteins, amino acids, RNA, and lipids are all grown in proportion. Since the membrane voltage also has been implicated in amino acid import [70, 71], it is likely a major global cell signal that influences cell growth and metabolism. Indeed, we observe that quiescent and senescent cells are depolarized, suggesting that membrane voltage reflects the cell metabolic state.

Our results show that at longer time scales, the membrane potential can be quantitatively modeled using the framework in Fig. 2. The overall potential is governed by the Poisson equation together with mobile ions and anionic lipids. Mobile ions can interact with anionic lipids, and their concentrations are described by a diffusion equation with fluxes from the cell boundaries. Ion fluxes from the cell boundaries are influenced by the membrane potential and also anionic lipids. The parameters of this system, i.e., lipid trafficking and conversion rates, ion channel fluxes, are further controlled by the actions of the cytoskeleton, transport motors, phosphorylation activity and signaling networks with multiple feedbacks. Therefore, a networked system of interactions is emerging that govern global cell properties such as, voltage, cell size, growth, and lipid metabolism.

Our findings suggest that membrane potential and anionic lipid trafficking may contribute to the establishment or maintenance of cell polarity and migration. Local voltage depolarization at protrusive regions, together with lateral enrichment of negatively charged lipids, is consistent with local protrusion formation and front–rear cell organization, although the precise causal relationships remain to be fully determined. This interpretation aligns with recent work showing that patterned modulation of membrane surface charge can trigger cell protrusions and stabilize cell polarity [72].

Our results demonstrates that cells are electromechanical objects. Cell mechanical activity, including cell spreading, substrate stiffness change, and external electric field exposure can alter membrane potential. The membrane potential acts as an integrator of mechanical, metabolic, and electrical information. Notably, membrane depolarization marks transitions into quiescent and senescent cell states, supporting a model where membrane potential serves as a biophysical indicator of cellular metabolic activity and growth status. Together, all of the results call for a broader model of membrane potential regulation-one that transcends the classical ion-centric view and incorporates lipid dynamics, cytoskeletal feedback, and mechanotransduction. Such an integrative framework may shed light on how cells coordinate membrane voltage with signaling pathways, polarity, and fate decisions across diverse physiological and pathological contexts.

## Materials and Methods

### Cell Culture

NIH 3T3 fibroblasts and HT1080 fibrosarcoma cells were obtained from ATCC and maintained in Dulbecco’s Modified Eagle Medium (DMEM; Corning), supplemented with 10% fetal bovine serum (FBS; Sigma) and 1% antibiotic solution containing 10,000 units/mL penicillin and 10,000 *μ*g/mL streptomycin (Gibco). All cell cultures and live-cell imaging experiments were performed at 37°C in a humidified incubator with 5% CO_2_.

### Osmotic Shock, Mitotic Swelling, Cell Cycle Voltage Measurement, and Pharmacological Inhibitor Experiments

For all experiments, 24-well glass-bottom plates were coated with 50 *μ*g/mL type I rat-tail collagen (Enzo) for 1 hour at 37°C. After incubation, plates were washed with the appropriate experimental media before cell seeding. Cells were seeded at a density of 5,000 cells/cm^2^ and cultured overnight in media containing 1 *μ*g/mL doxycycline. Two hours before imaging, the doxycycline-containing media was removed, cells were washed once with PBS, and replaced with fresh cell culture media.

For the osmotic shock experiments, cells were pre-incubated overnight in isotonic solution (329 mOsm)[73], prepared by mixing 50% Dulbecco’s phosphate-buffered saline (without calcium and magnesium, Sigma-Aldrich) with 50% culture media. Hypotonic solution (175 mOsm) was prepared using 50% ultra-pure water and 50% culture media. Osmolality was measured using an Advanced Instruments Model 3320 osmometer.

For mitotic swelling and cell cycle voltage experiments, cells were imaged continuously for 30 hours using time-lapse microscopy. For mitotic swelling analysis, data were extracted from the 30-minute window preceding cell division. For cell cycle voltage analysis, only cells that completed a full cell cycle from one division to the next were analyzed.

For pharmacological inhibition experiments, cells were treated with the following compounds (from Tocris unless otherwise specified): DMSO (vehicle control, 0.2%, Invitrogen), Latrunculin A (2 *μ*M), Gramicidin (20 nM), Valinomycin (10 *μ*M), NSC668394 (20 *μ*M, MilliporeSigma), Ouabain (100 *μ*M), EIPA (40 *μ*M), Bumetanide (40 *μ*M), 4-Aminopyridine (4-AP, 4 mM), BAPTA Tetrapotassium Salt (1.2 mM, Invitrogen), BAPTA-AM (10 *μ*M, Invitrogen), Syrosingopine (10 *μ*M), DIDS (40 *μ*M), Dynasore (80 *μ*M), MyoVin (80 *μ*M), and LY294002 (20 *μ*M). For all membrane potential measurements, cells were treated for 2 hours unless otherwise stated: EIPA was applied for 1 hour, and ouabain for 4 hours.

### Validation of JEDI-2P/cyOFP1 Ratiometric Voltage Indicator

To assess whether the JEDI-2P/cyOFP1 fluorescence ratio remains stable independent of expression levels, we cultured cells in 24-well plates and induced JEDI-2P expression using doxycycline overnight. Following induction, doxycycline was removed, and cells were imaged every 2 hours over 12 hours. At each time point, we calculated the cpGFP/cyOFP1 ratio across cells to evaluate its stability over time. Despite gradual decreases in total fluorescence intensity due to protein degradation or dilution, the ratiometric signal remained consistent throughout the time course (Fig. 1A, B, C, D, E, F), indicating that the JEDI-2P/cyOFP1 ratio is robust against fluctuations in expression level for both 3T3 and HT1080 cells.

In addition, we observed that JEDI-2P was localized not only to the plasma membrane but also to intracellular organelles, including the endoplasmic reticulum (ER), Golgi apparatus, and endosomes. Shortly after doxycycline withdrawal (1-4 hours), strong cpGFP fluorescence was detected in the ER and Golgi. As the withdrawal period extended (≥ 6 hours), JEDI-2P was gradually exported from these organelles, with residual cyOFP1 fluorescence persisting primarily in endosomes (Fig. S1-G, Fig. 1B). These observations confirm the intracellular trafficking dynamics of JEDI-2P and further support the stability of the ratiometric readout across subcellular compartments.

### 3D Collagen gel matrix

Cell-impregnated 3D collagen matrices were prepared by mixing cells suspended in culture medium with 10X reconstitution buffer (1:1, v/v), and then combining the mixture with soluble rat tail type I collagen in acetic acid (BD Biosciences) to achieve a final collagen concentration of 2 mg/mL. To adjust the pH to 7.0, 10–20 *μ*L of 1 M NaOH was added. A volume of 150 *μ*L of the final mixture was transferred into each well of a 24-well plate and incubated at 37°C in a humidified incubator with 5% CO_2_ for 1 hour to allow gel solidification. Subsequently, 500 *μ*L of cell culture medium containing 1 *μ*/mL doxycycline was added to each well. Prior to imaging, the doxycycline-containing medium was removed, the collagen gels were washed once with PBS, and then replaced with fresh cell culture medium. A collagen concentration of 2 mg/mL was selected to ensure that the matrix pore size (*<*1 *μ*m) was significantly smaller than the cell body and nucleus. We verified that cells continued to proliferate normally within the 3D matrix for over 48 hours.

### Cell Spreading Experiment

For the spreading experiments, cells were treated with 1 *μ*g/mL doxycycline overnight. Glass-bottom 24-well plates were coated with 50 Âμg/mL type I rat-tail collagen (Enzo) for 1 hour at 37°C, followed by a wash with isotonic media. Each well was filled with 1 mL of isotonic media prior to cell seeding. Before the experiment, cells were concentrated to approximately 1-2 million cells/mL, and 10 *μ*L of the suspension was added to each well. The plates were then immediately placed on the microscope, and Z-stack images were acquired to capture cells before spreading began.

### Confocal Microscopy

Confocal imaging was performed using a Zeiss LSM 800 confocal microscope equipped with either a 20*×* air 0.8-NA objective or a 63*×* oil-immersion 1.2-NA objective. All live-cell imaging was conducted using a stage-top incubator (Pecon) with a CO_2_ Module S and TempModule S (Zeiss), maintained at 37°C with 5% CO_2_. Immunofluorescence imaging was conducted at room temperature without CO_2_. Image acquisition was performed using ZEN 2.6 or 3.6 software (Zeiss), and image analysis was carried out using custom MATLAB (MathWorks) scripts or ImageJ.

For membrane potential measurements using the ratiometric indicator JEDI-2P-cyOFP1, cells were excited at 488 nm; cpGFP emission was collected from 400-570 nm and cyOFP1 from 570-700 nm. For anionic lipid measurements using the ratiometric indicators AktPH-CaaX and Lact-C2-CaaX, CaaX was excited at 488 nm and detected at 400-570 nm, while AktPH and Lact-C2 were excited at 561 nm with emission collected from 570-700 nm. For experiments involving significant changes in cell shape (e.g., EIPA treatment, LatA treatment, and spreading assays), time-lapse Z-stack imaging was performed using a 1 *μ*m step size over a total imaging depth of 20 *μ*m.

### Stable Cell Line Generation, Lentivirus Preparation, Transduction, and Transfection

PiggyBac vectors were generated by cloning PCR-amplified constructs into the PiggyBac vector backbone (Addgene, a gift from Volker Busskamp) using HiFi DNA Assembly (New England Biolabs). Wild-type 3T3 cells were transfected with a 9:1 ratio of PiggyBac plasmid to Super PiggyBac transposase plasmid using Lipofectamine 3000 (Thermo Fisher Scientific), following the manufacturer’s instructions. To establish a stably integrated transgenic cell line, culture medium was supplemented with *μ*g/mL puromycin (Gibco) beginning one day post-transfection. Puromycin-containing medium was refreshed every 2-3 days until cells reached confluence and were subsequently cryopreserved.

For lentivirus production, HEK 293T/17 cells were co-transfected with psPAX2, VSV-G, and the lentiviral transfer plasmid using standard protocols. Forty-eight hours after transfection, viral supernatants were collected and concentrated via centrifugation. Wild-type 3T3 cells at 60-80 *μ*g/mL Polybrene (Millipore Sigma) to enhance transduction efficiency.

### Live Cell Reporters

To monitor PIP3 dynamics, pcDNA3.1-AktPH-mCherry (Addgene 67301) was used for transient transfection. 3T3 cells at 60–80% confluency were transfected using Lipofectamine 3000 (Thermo Fisher Scientific) following the manufacturer’s protocol. For cell membrane labeling and intracellular pH measurement, pHR GFP-CaaX (Addgene 113020) and FUGW-pHRed (Addgene 65742) were delivered via lentiviral transduction. Lentivirus production and transduction protocols are described in the previous section, and successfully transduced cells were sorted by flow cytometry.

To visualize phosphatidylserine (PS), mRFP-Lact-C2 (Addgene 74061) was used. For membrane potential measurements, the genetically encoded voltage indicator 2801-pcaggs-jedi2p-cyofp1 (a gift from Francois St-Pierre’s lab) was employed. Both the PS and voltage reporters were stably integrated into cells using the PiggyBac transposon system, as previously described.

### Cell Voltage, PIP_3_, and PS Data Analysis

A consistent analysis pipeline was applied to all membrane potential (JEDI), PIP3, and PS datasets to ensure reproducibility. For each cell, a rectangular region was cropped, and the outer area of the cropped image was used to calculate the background intensity for each fluorescence channel, which was subtracted to improve the signal-to-noise ratio. A binary mask was generated to isolate the cell, and the cell boundary was defined as the outermost two-pixel layers of the mask. The boundary positions were clustered into 100 groups using the k-means algorithm and ordered sequentially in a clockwise direction to ensure continuity. To calculate the overall ratio for each cell, the mean of the intensity ratio values across all 100 boundary groups was computed after background subtraction. For distribution analysis, two points along the boundary were manually selected to define the region of interest. The boundary groups closest to the start and end points were identified, and a ratio curve was plotted from the first to the last group, with each point representing the average ratio of one group. To assess signal uniformity, we calculated the difference between the maximum and minimum values of the 100-group ratio curve, normalized by the mean ratio value, i.e., Δ*R/* ⟨*R*⟩ = (*R*_max_ −*R*_min_)*/* ⟨*R*⟩. For Z-stack time-lapse images, the focal plane with the highest cyOFP1 intensity at each time point was identified and treated as the base layer. All analyses described above were performed on this selected layer to ensure consistency across time points.

### FRAP Experiment and Membrane Fusion Rate Calculation

Fluorescence recovery after photobleaching (FRAP) experiments were performed using a Zeiss LSM800 confocal microscope equipped with a 63*×* oil-immersion objective and a laser bleaching module. 3T3 cells stably expressing the phosphatidylserine (PS) biosensor mRFP-Lact-C2 (Addgene 74061) were seeded into 24-well plates and incubated overnight prior to imaging. To assess membrane dynamics at different cell regions, rectangular regions of interest (ROIs) were manually positioned at the lateral and protrusion regions of the membrane. Each ROI was photobleached using a 561 nm laser at 50% intensity for 100 iterations. Immediately following bleaching, time-lapse imaging was conducted for 5 minutes at a 10-second frame interval to monitor fluorescence recovery. The average PS intensity within the bleached ROI was quantified at each time point, and the recovery curve was fitted using an exponential model to evaluate membrane fusion rates.

### PDMS Substrate Fabrication and Experiment Preparation

Soft PDMS substrates with defined stiffnesses were fabricated to modulate substrate mechanics. For 5 kPa substrates, Sylgard 527 Part A and Part B (Dow) were mixed at a 1:1 weight ratio. For 400Pa substrates, QGel 920 Part A and Part B (Grace Bio-Labs) were mixed at a 1:1 ratio, as previously described[74]. To prepare 15 kPa and 130 kPa substrates, 2% and 20% Sylgard 184 (10:1 base-tocuring-agent ratio) were added to Sylgard 527 (1:1 base-to-curing-agent ratio), respectively, following published protocols[75]. All PDMS mixtures were vacuum-degassed for approximately 5 minutes to remove air bubbles and spin-coated onto 35mm or 50mm glass-bottom dishes at 1,000 rpm for 60 seconds. The substrates were cured at room temperature for 24 hours. The final Young’s modulus of each substrate was confirmed using a rheometer. Prior to cell seeding, all substrates were sterilized under UV light for at least 15 minutes. The glass-bottom dishes were then coated with 50 *μ*g/mL type I rat-tail collagen (Enzo) for 1 hour at 37°C, followed by thorough washing with experimental media. Cells were seeded at a density of 5,000 cells/cm^2^, and 1 *μ*g/mL doxycycline was added to induce expression of transgenes. After overnight culture, doxycycline-containing media were removed, cells were washed once with PBS, and replaced with fresh culture media 2 hours before imaging.

### Mechanical Stretch

For mechanical stretch experiments, 3T3 cells were seeded into elastic PDMS chambers (STB-CH-4W, STREX) at a density of 5,000 cells/cm^2^ in complete culture media. Prior to seeding, the chambers were coated with 200 *μ*g/mL type I collagen (Enzo) for 4 hours at 37°C to promote cell adhesion. A single uniaxial stretch of 20% strain was applied using a motorized stretching device (STB-100, STREX) for downstream live-cell imaging and analysis.

### Immunofluorescence

Cells were seeded into collagen-I coated (50 Âμg/mL for 1 h in 37°C) glass bottom 24 well plates at a density of 5000 cells/cm^2^ overnight. For NHE1 staining, cells were fixed with 4% paraformaldehyde (ThermoFisher Scientific) at different time points as indicated. After washing 3X with PBS, cells were permeabilized in 0.1% Triton X-100 for 15 min. Then, cells were washed 3X in PBS and blocked for 1 h at RT with 5% BSA solution containing 5% normal goat serum (Cell Signaling) and 0.1% Triton X-100. Next, cells were incubated at 4°C with primary antibody overnight. After washing 3X with PBS, samples were incubated for 1 h at RT with a secondary antibody. Primary antibody: anti-NHE1 (Invitrogen; PA5-116471; 1:100). Secondary antibody: Alexa Fluor 488 goat anti-rabbit immunoglobulin G (IgG) H+L, (Invitrogen; A11008; 1:200). For actin staining, Alexa Fluor 647 Phalloidin (Invitrogen; A22287; 1:100) were added for 1h at RT.

### Intracellular pH Measurement

For intracellular pH measurements, cells expressing the ratiometric pH indicator pHRed were prepared as described previously. Imaging was performed using a confocal microscope with excitation at 405 nm and 561 nm, and emission collected above 550 nm. The intracellular pH was inversely correlated with the fluorescence intensity ratio (*I*_561_*/I*_405_). Cells were seeded into collagen-I-coated glass-bottom 24-well plates (50 *μ*g/mL, 1h at 37°C) at a density of 5,000 cells/cm^2^ in complete media. Before imaging, cells were allowed to equilibrate on the microscope stage for at least 15 minutes in a humidified 37°C, 5% CO_2_ environment. Image analysis was performed using a pipeline consisting of cell masking, tracking, and background subtraction. The *I*_561_*/I*_405_ ratio was calculated for each cell to evaluate relative pH.

To obtain absolute pH values, cells were calibrated using the Intracellular pH Calibration Buffer Kit (Invitrogen, P35379) following the manufacturer’s instructions. Briefly, pHRed-expressing cells were incubated for at least 5 minutes in a calibration buffer containing 10 *μ*M valinomycin and 10 *μ*M nigericin at defined pH values of 5.5, 6.7, and 7.5. A linear calibration curve was generated from the measured fluorescence ratios. Experimental pH values for individual cells were obtained by fitting their fluorescence ratios to the curve. Relative intracellular pH change (ΔpH_i_) was calculated by comparing pH_i_ values before and after treatment or osmotic shock for each cell.

### Cell Cycle Arrest and Senescence Cell Induction

To induce cell cycle arrest at the late G1 phase, 3T3 cells were cultured in a medium containing 0.5% fetal bovine serum (FBS) for 24 hours to promote quiescence. For HT1080 cells, cell cycle arrest was induced by treating the cells with 10 *μ*M Palbociclib, a selective CDK4/6 inhibitor, for 24 hours prior to the experiment. Palbociclib was maintained in the culture medium throughout the imaging session.

To induce cellular senescence, cells at approximately 50% confluence were treated with 250 nM doxorubicin (SelleckChem, S1208) for 24 hours. Following treatment, cells were washed once with PBS and returned to standard culture media. The cells were then incubated under normal conditions for 7 days to allow full development of the senescent phenotype.

### Fitting the Point Spread Function (PSF), Applying Deconvolution, and Validation

To determine the point spread function (PSF) of our imaging system, we used 0.2 *μ*m red (FluoSpheres 633/660) and green (FluoSpheres 505/515) fluorescent beads (Invitrogen). The beads were diluted by mixing 5 *μ*L of stock solution into 2 mL of ethanol. Coverslips were cleaned with 100% isopropanol (IPA), and 10 *μ*L of the diluted bead solution was pipetted onto fluor dishes and allowed to dry in the dark. After drying, 10 *μ*L of non-hardening mounting medium was applied, and a coverslip was carefully placed on top, minimizing air bubbles using a cotton swab. Z-stack images were acquired using a Zeiss LSM800 confocal microscope with a 20× objective at 0.2 *μ*m intervals. To extract the PSF, bead images were processed using a custom MATLAB script. The brightest 4 × 4 pixel regions from each bead were selected to obtain intensity profiles along the z-axis, and a one-dimensional Gaussian function was fitted to determine the PSF [76]. The fitted full width at half maximum (FWHM) was 0.83 *μ*m for green fluorescence and 0.92 *μ*m for red fluorescence (Fig. S2A). The Richardson-Lucy deconvolution algorithm was implemented in MATLAB and applied to each z-stack for both fluorescence channels.

To evaluate whether deconvolution altered our core findings, we applied it to JEDI-expressing 3T3 cells and reanalyzed membrane potential distributions at the leading, trailing, and lateral edges. The results before and after deconvolution were nearly identical (Fig. S2B, C, D, E), confirming the stability of our original observations. Although the difference between leading and trailing regions was not statistically significant, the trailing edge consistently showed slightly higher voltage, as observed in pre-deconvolution analysis. We further validated the deconvolution method on datasets with large polarity changes, including cells treated with EIPA and Latrunculin A (LatA), as well as during cell spreading conditions where PSF distortion could be most pronounced. Even after deconvolution, both EIPA and LatA treatments caused hyperpolarization and increased membrane potential uniformity (Fig. S5E, F, G), fully consistent with the original results. In spreading assays, deconvolved images showed continued depolarization post-spreading and preserved the negative correlation between JEDI ratio and cell area (Fig. S5F, G, H). Finally, we applied deconvolution to datasets measuring PIP3 and PS distributions. Post-processing results showed lateral enrichment of both anionic lipids remained unchanged (Fig. S3B-G), indicating that our measurements of lipid localization were not artifacts of PSF but reflected true biological distributions.

### Electrotaxis experiment

To study electrotaxis, we fabricated a PDMS-based microfluidic device with a 50 *μ*m high seeding area using photolithography, followed by a standard replica molding technique, as previously described [77, 78]. A 50 *μ*m thick layer of SU-8 was spin-coated onto a silicon wafer, cross-linked, and patterned with UV light through a photomask to define the seeding and electrode areas. PDMS and the curing agent were mixed at a 10:1 ratio, poured onto the patterned wafer, and baked at 85°C for 120 minutes to cure. The final device contained six seeding wells and two electrode wells, spaced 1 cm apart. Before cell seeding, devices were coated with collagen I (20 *μ*g/mL) (Thermo Fisher Scientific) and incubated at 37°C with 5% CO_2_ for at least 1 hour to enhance cell adhesion. For electrode fabrication, Ag/AgCl electrodes (EP2, WPI), 2 mm in diameter and 4 mm in length, were cleaned and immersed in cell culture-grade agarose (Thermo Fisher Scientific) within a 300 *μ*L Eppendorf tube. After cooling to room temperature, the gelled agarose was trimmed, and electrodes were stored in PBS (Gibco) until use. During electrotaxis experiments, electrodes were positioned on either side of the seeding area, and a 3 V/cm DC voltage was applied using a programmable potentiostat (Interface 1010E, Gamry Instruments). Live-cell imaging was performed using a Nikon A1 or AXR confocal microscope with a Plan Apo 20× /0.75 NA objective (A1) or a Plan Apo 20 × /0.8 NA objective (AXR). Cells were maintained at 37°C and 5% CO_2_ in a stage-top incubator (Tokai Hit), with both confocal systems equipped with a temperature-controlled cage to ensure stable imaging conditions.

To quantify the fraction of cells that respond to electric field stimulation, we compared cell morphology before and 10 minutes after applying a 3 V/cm DC electric field. A cell was classified as voltage-sensitive if a new protrusion was formed in the direction of the applied field. Cells that showed no change in morphology or generated protrusions in directions unrelated to the field orientation were classified as not voltage-sensitive.

### Statistical Analysis

For time series plots, error bars represent the mean and the standard error of the mean (SEM) from at least three biological replicates. The number of cells analyzed for each condition is reported in the corresponding figure panels or captions. For scatter plots, error bars represent the mean and the standard deviation (SD) from at least three biological replicates.

The normality of the data was assessed using the D’Agostino-Pearson omnibus normality test. For datasets that did not follow a normal distribution, statistical comparisons between two conditions were performed using the nonparametric Mann-Whitney U test. Comparisons involving more than two groups were analyzed using the Kruskal-Wallis test followed by Dunn’s multiple-comparisons test. All statistical analyses were performed using GraphPad Prism 10 (GraphPad Software). Statistical significance was defined as *p <* 0.05. Significance levels are indicated as follows: ns (not significant), \**p<* 0.05, \*\**p<* 0.01, \*\*\**p<* 0.001, and \*\*\*\**p<* 0.0001.

---

**Figure S1.**
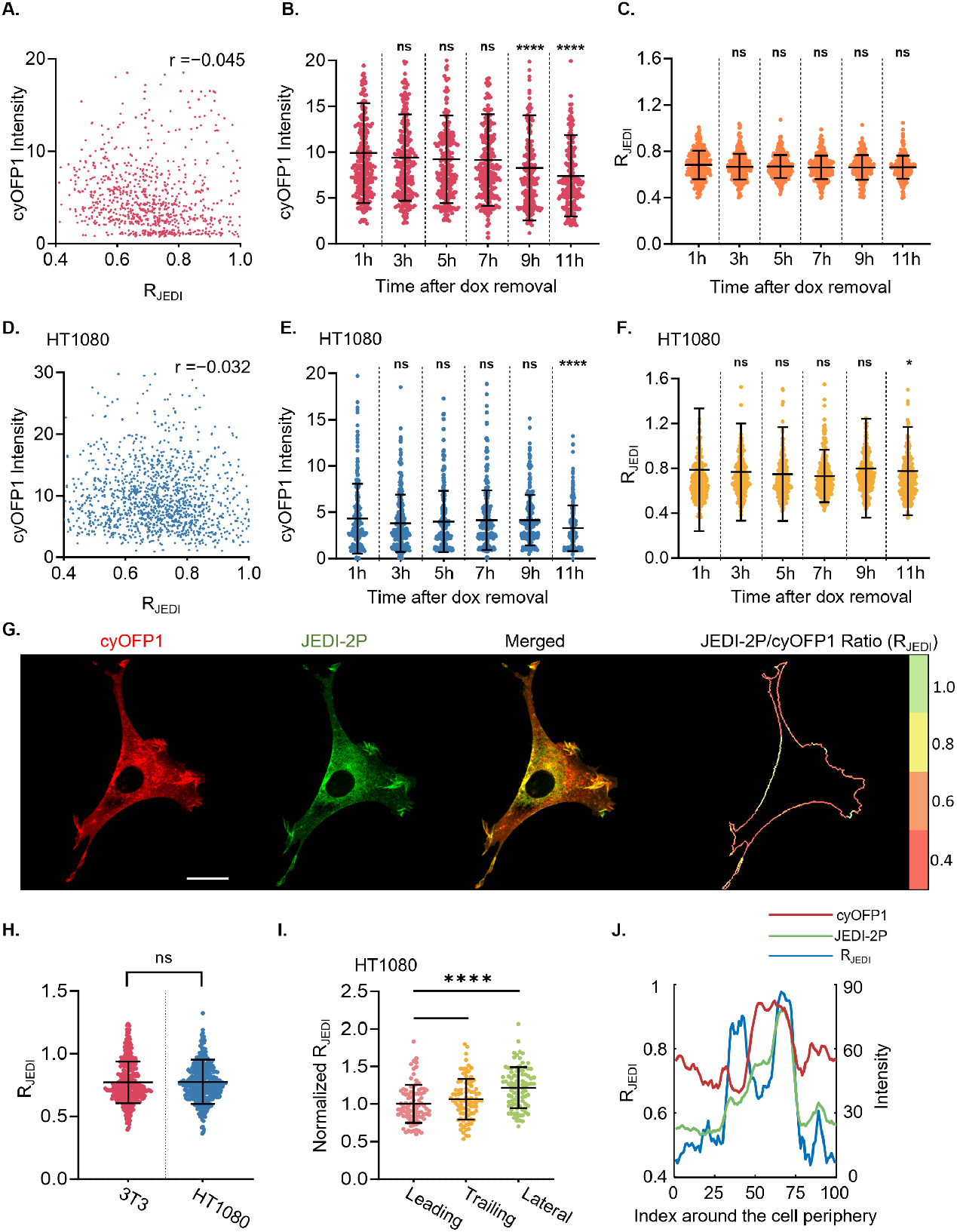
JEDI-2P expression validation and membrane potential comparison between 3T3 and HT1080 cells. **(A, D)** Scatter plots of R_JEDI_ versus cyOFP1 intensity in 3T3 (A) and HT1080 (D) cells. Pearson correlation coefficients (*r*) are shown. *n*_3T3_ = 944, *n*_HT1080_ = 1371. **(B, E)** Time-course of cyOFP1 intensity after doxycycline (dox) removal in 3T3 (B) and HT1080 (E) cells. *n*_3T3_ = 248, 259, 259, 217, 210, *n*_HT1080_ = 244, 316, 259, 259, 253, 238. **(C, F)** Time-course of R_JEDI_ in 3T3 (C) and HT1080 (F) following dox removal; sample sizes as in (B) and (E). **(G)** Representative confocal images of 3T3 cells expressing JEDI-2PcyOFP1 within 4h dox removal, with R_JEDI_ mapped along the cell boundary. **(H)** Comparison of mean R_JEDI_ between 3T3 and HT1080 cells. *n*_3T3_ = 321, *n*_HT1080_ = 362. **(I)** Normalized R_JEDI_ in HT1080 cells across leading, trailing, and lateral regions. *n*_HT1080_ = 91. **(J)** Distribution of R_JEDI_, absolute JEDI-2P, and cyOFP1 intensity along the cell boundary on a 2D glass substrate (from 1B). Blue: R_JEDI_ (left y-axis); red: cyOFP1 (right y-axis); green: JEDI-2P (right y-axis). **(B, C, E, F, H, I)** Error bars represent standard deviation unless noted. (B, C, E, F) Kruskal-Wallis tests with Dunn’s multiple-comparisons were performed relative to the 1h condition. (H) Error bars indicate SEM. Mann-Whitney tests were used for two-condition comparisons. (I) Kruskal-Wallis with Dunn’s multiple-comparisons was used relative to the leading-edge region. **(G)** Scale bar: 20 *μ*m.

**Figure S2.**
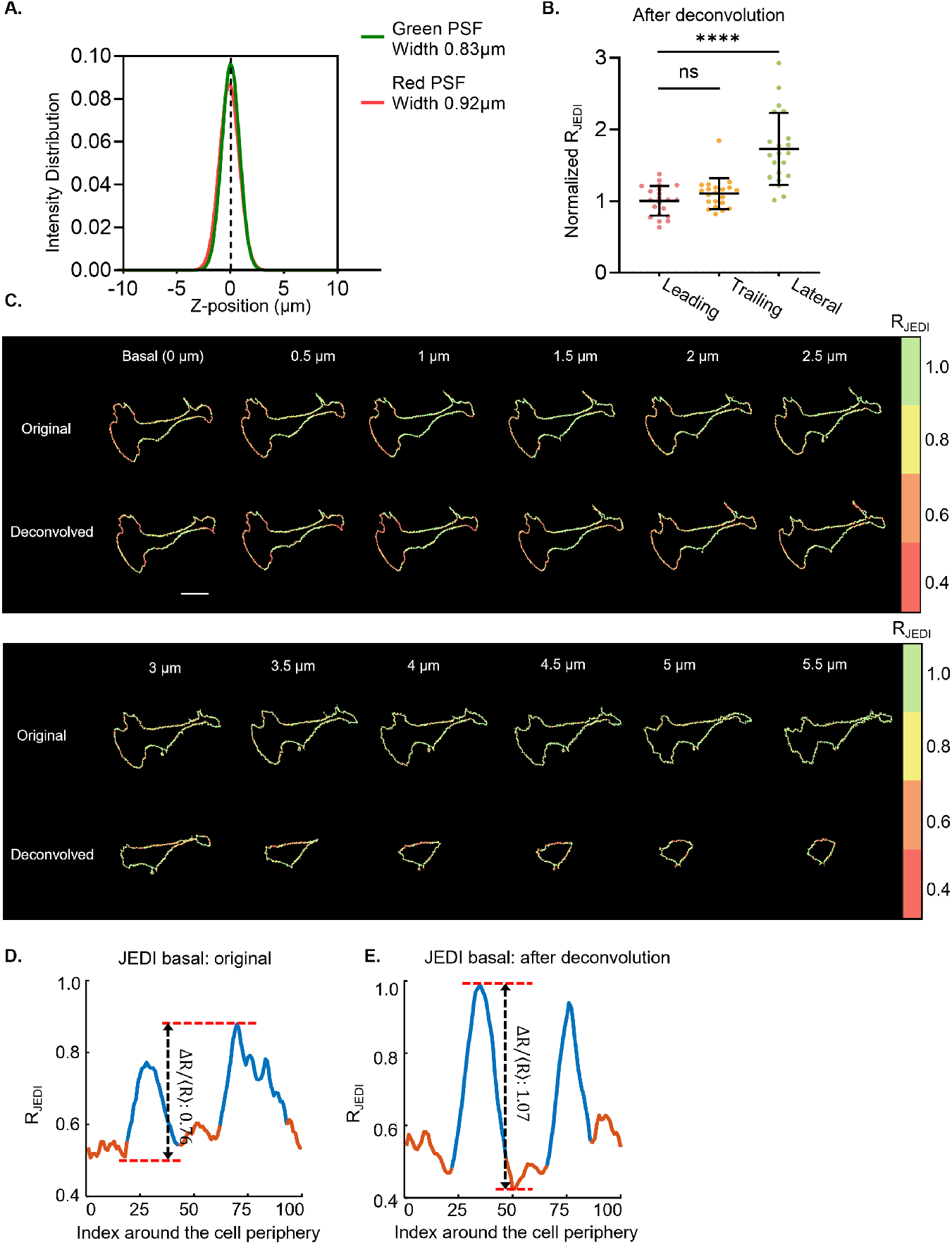
Deconvolution enhances JEDI-2P signal resolution while perserving membrane potential distribution. **(A)** Point spread function (PSF) measurement for green (488 nm excitation, 500-570 nm emission) and red (561 nm excitation, 570-700 nm emission) channels, with respective widths of 0.83 *μ*m and 0.92 *μ*m. **(B)** Normalized *R*_JEDI_ across membrane regions after deconvolution, showing trends consistent with undeconvolved data (Fig.1D). *n*_3T3_ = 21. **(C)** Representative confocal images of an NIH 3T3 cell expressing JEDI-2P-cyOFP1, showing optical slices (0-5.5 *μ*m) before and after deconvolution, corresponding to the 3D reconstruction in Fig. 1G. **(D, E)** *R*_JEDI_ intensity profiles along the cell boundary before (D) and after (E) deconvolution. **(B)** Error bars indicate the standard deviation. Kruskal-Wallis tests followed by Dunn’s multiple-comparisons test were performed relative to the leading-edge region. **(C)** Scale bar: 20 *μ*m.

**Figure S3.**
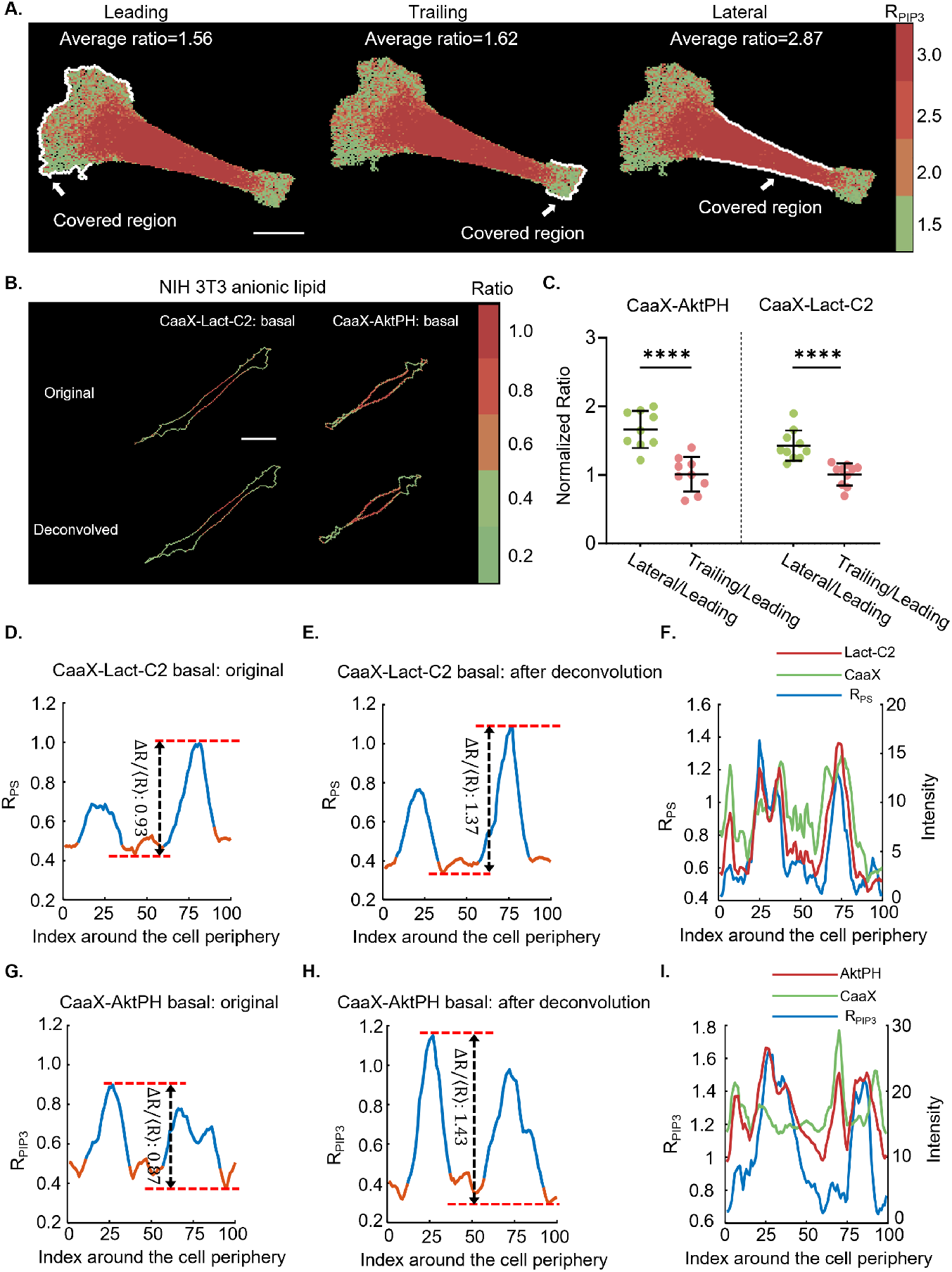
PIP_3_ spatial analysis and deconvolution confirm anionic lipid distribution in 3T3 cells. **(A)** Illustration of how *R*_PIP3_ is calculated in the leading, trailing, and lateral regions by averaging the intensity values within each area. The same approach is used for *R*_PS_ and *R*_JEDI_. **(B)** Representative confocal images of NIH 3T3 cells expressing AktPH-CaaX and Lact-C2-CaaX before and after deconvolution. **(C)** Normalized *R*_PIP3_ and *R*_PS_ across membrane regions after deconvolution, showing trends consistent with predeconvolution (Fig. 2G). *n*_PIP3_ = 9, *n*_PS_ = 10. **(D, E)** *R*_PS_ intensity profiles along the cell boundary before (C) and after (D) deconvolution. **(F)** Distribution of R_PS_, absolute CaaX, and Lact-C2 intensity along the cell boundary (from 2D). Blue: R_PS_ (left y-axis); red: Lact-C2(right y-axis); green: Caax (right y-axis). **(G**,**H)** *R*_PIP3_ intensity distribution along the cell boundary before (G) and after (H) deconvolution. **(I)** Distribution of R_PIP3_, absolute CaaX, and AktPH intensity along the cell boundary (from 2C). Blue: R_PIP3_ (left y-axis); red: AktPH(right y-axis); green: Caax (right y-axis). **(C)** Error bars indicate the standard deviation. MannWhitney tests were used for two-condition comparisons. **(A, B)** Scale bar: 20 *μ*m.

**Figure S4.**
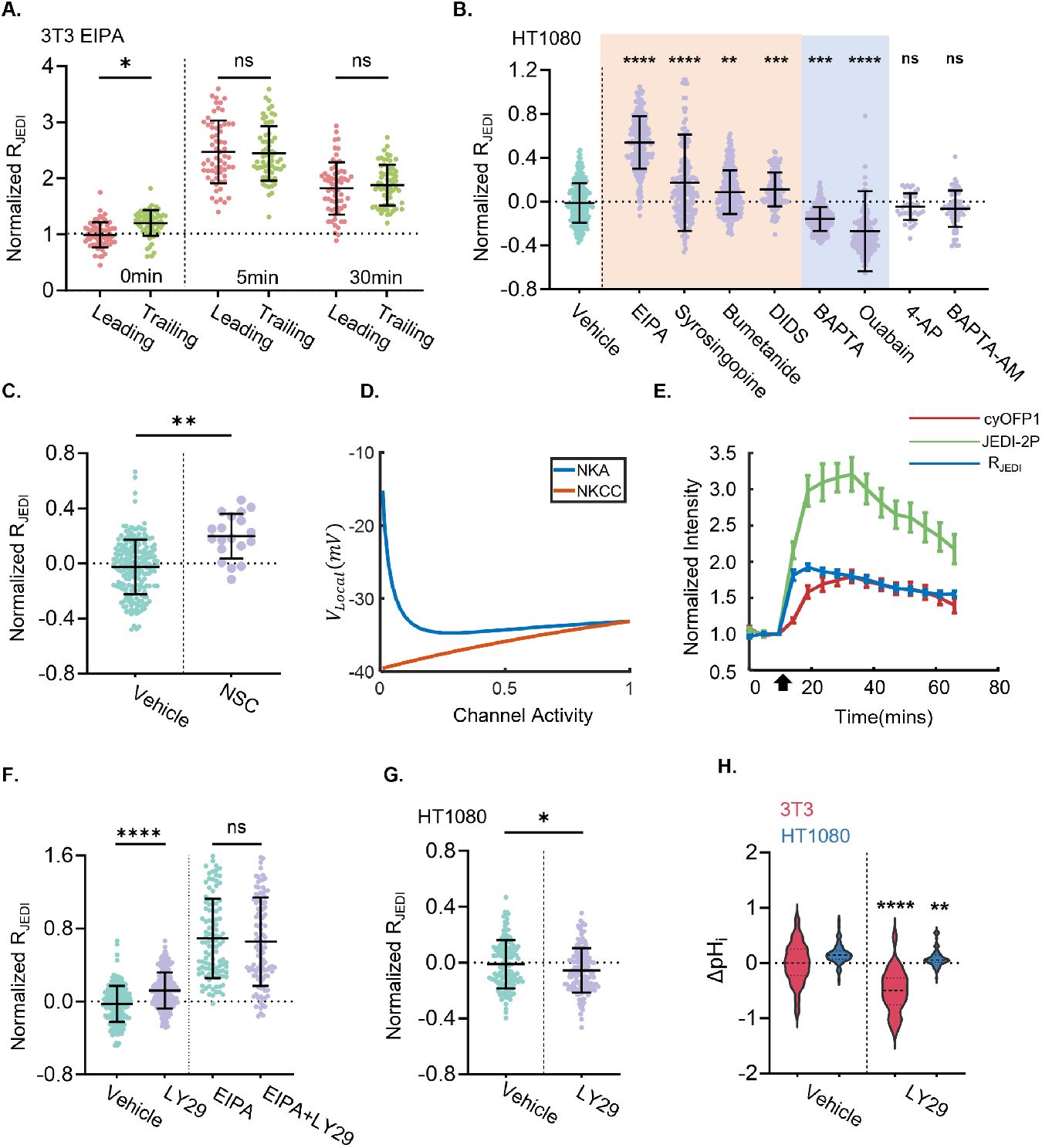
Effects of ion transporter inhibition and PI3K signaling on membrane potential. **(A)** Time-dependent *R*_JEDI_ changes in 3T3 cells showing the leading-to-trailing ratio after NHE inhibition. *n*_3T3_ = 62. **(B)** Normalized *R*_JEDI_ in HT1080 cells treated with various ion transport inhibitors (vehicle, EIPA, Syrosingopine, Bumetanide, DIDS, BAPTA, Ouabain, 4-aminopyridine, and BAPTA-AM) at the same doses as in Fig. 3A,E. *n*_HT1080_ = 279, 264, 166, 286, 143, 275, 166, 65, 39. **(C)** Normalized *R*_JEDI_ in 3T3 cells treated with vehicle or NSC (20 *μ*M, Ezrin protein inhibitor). *n*_3T3_ = 200, 19. **(D)** Model predictions of voltage *V*_Local_ under varying NKA and NKCC activity with low potassium permeability. **(E)** Time-lapse normalized intensity of *R*_JEDI_, JEDI-2P, and cyOFP1 in 3T3 cells following EIPA treatment (*n*_3T3_ = 200). **(F)** Normalized *R*_JEDI_ in 3T3 cells treated with vehicle, LY294002 (LY29, 20 *μ*M; PI3K inhibitor), EIPA, or EIPA + LY29. *n*_3T3_ = 200, 253, 108, 108. **(G)** Normalized *R*_JEDI_ in HT1080 cells treated with vehicle and LY29. *n*_HT1080_ = 132, 136. **(H)** Relative intracellular pH change (ΔpH_*i*_) in 3T3 and HT1080 cells treated with vehicle and LY29. *n*_3T3_ = 60, 28. *n*_HT1080_ = 87, 63. **(A, C, F, G, H)** Error bars represent standard deviation. ((A, C, F, G, H)) Mann-Whitney tests were used for two-group comparisons. (B) Kruskal-Wallis tests with Dunn’s multiple-comparisons test were performed against vehicle control. **(E)** Error bars represent the standard error of the mean (SEM).

**Figure S5.**
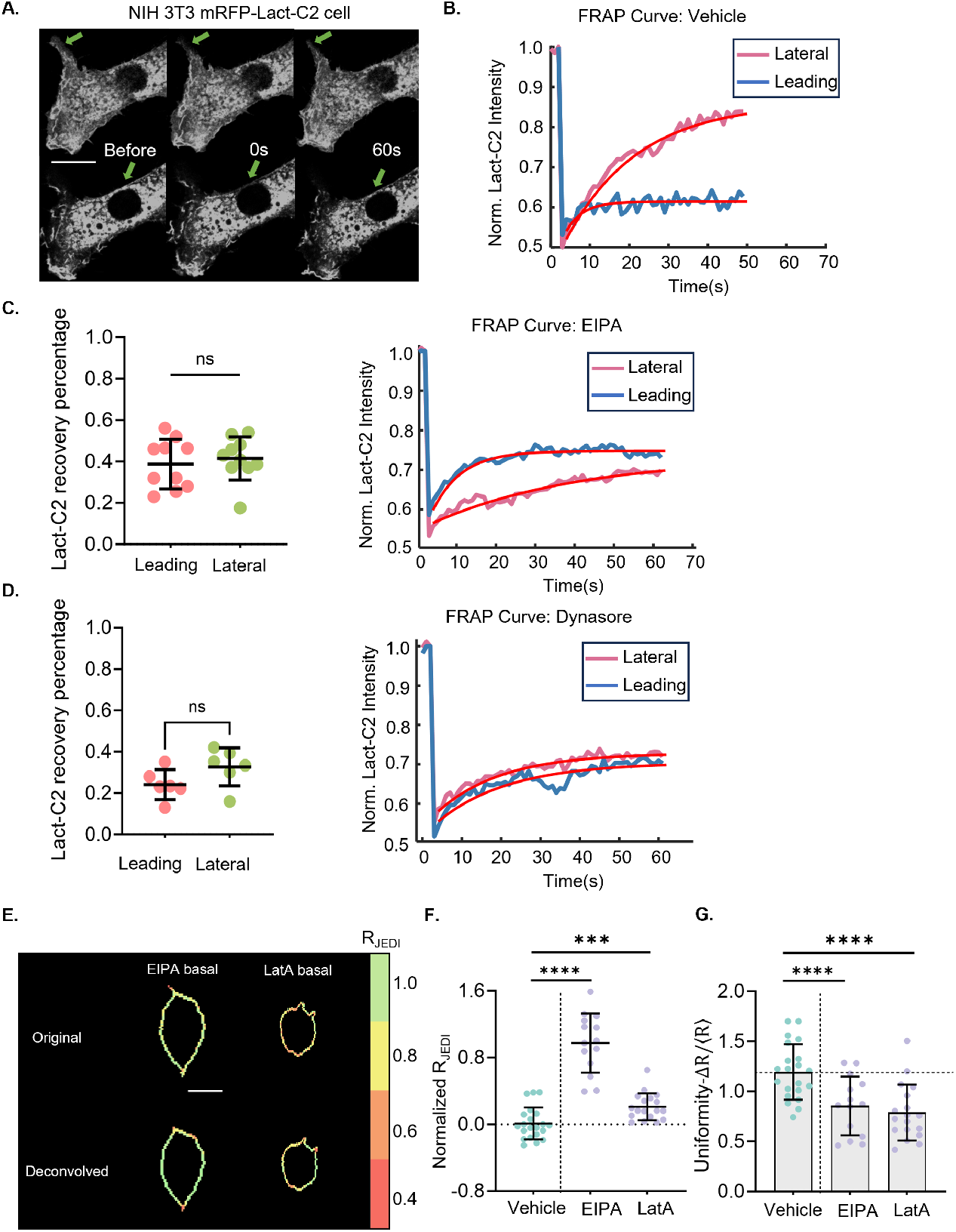
FRAP analysis of membrane trafficking and JEDI distribution after endocytosis inhibition. **(A)** Representative confocal images of a 3T3 cell before and after photobleaching (0s and 60s), with green arrows marking bleached membrane regions. **(B)** Normalized Lact-C2 intensity recovery curves for lateral and protrusion membrane regions. **(C)** Quantification of the final Lact-C2 recovery percentage in leading and lateral regions, and recovery curves during EIPA treatment. *n*_3T3_ = 10. **(D)** Quantification of final Lact-C2 recovery percentage in the leading and lateral regions, and recovery curves during Dynasore treatment. *n*_3T3_ = 6. **(E)** Representative confocal images of 3T3 cells expressing JEDI-2P-cyOFP1 before and after deconvolution following EIPA and LatA treatment. **(F)** Normalized *R*_JEDI_ in 3T3 cells treated with EIPA and LatA following deconvolution, showing trends consistent with pre-deconvolution (EIPA in Fig. 3A, LatA in Fig. S8C). *n*_3T3_ = 21, 14, 18. **(G)** Quantification of voltage uniformity in 3T3 cells treated with EIPA and LatA following deconvolution, showing trends consistent with pre-deconvolution (Fig. 4B). **(C, D, F, G)** Error bars indicate the standard deviation. (C, D) Mann-Whitney tests were used for two-group comparisons. (F, G) Kruskal-Wallis tests with Dunn’s multiple-comparisons test were performed against vehicle control. **(A, E)** Scale bar: 20 *μ*m.

**Figure S6.**
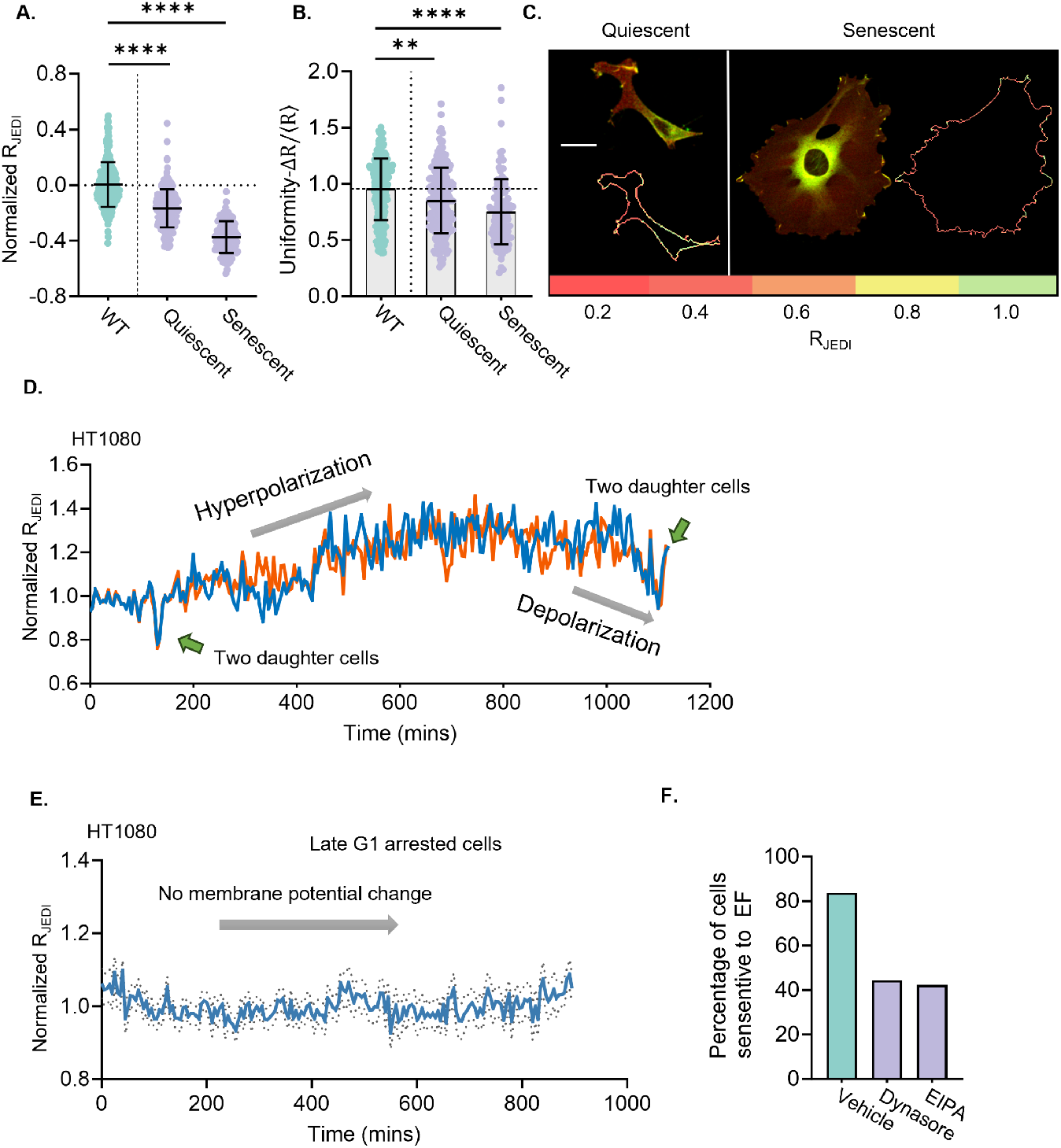
Membrane potential dynamics during cell cycle progression and EF response after endocytosis inhibition. **(A)** Normalized *R*_JEDI_ in 3T3 cells induced into quiescence (0.5% FBS overnight) or senescence (doxorubicin 250 nM for 24 h, followed by 7 days of culture). *n*_3T3_ = 214, 207, 110 **(B)** Quantification of voltage uniformity in quiescent and senescent 3T3. **(C)** Representative confocal images of 3T3 cells expressing JEDI-2P-cyOFP1 under quiescent and senescent conditions, with *R*_JEDI_ mapped along the cell boundary. **(D)** Representative trajectories of normalized *R*_JEDI_ in HT1080 cells over a full cell cycle, showing hyperpolarization and depolarization events across different cell cycle phases. **(E)** Normalized *R*_JEDI_ in HT1080 cells arrested in late G1 using palbociclib (10 *μ*M, CDK4/6 inhibitor), showing stable membrane potential over time. *n*_HT1080_ = 18. **(F)** Percentage of 3T3 cells responding to applied electric field during endocytosis inhibition. **(A, B, E)** Error bars indicate the standard deviation. Kruskal-Wallis tests with Dunn’s multiple-comparisons versus wild type (WT) controls. **(C)** Scale bar: 20 *μ*m.

**Figure S7.**
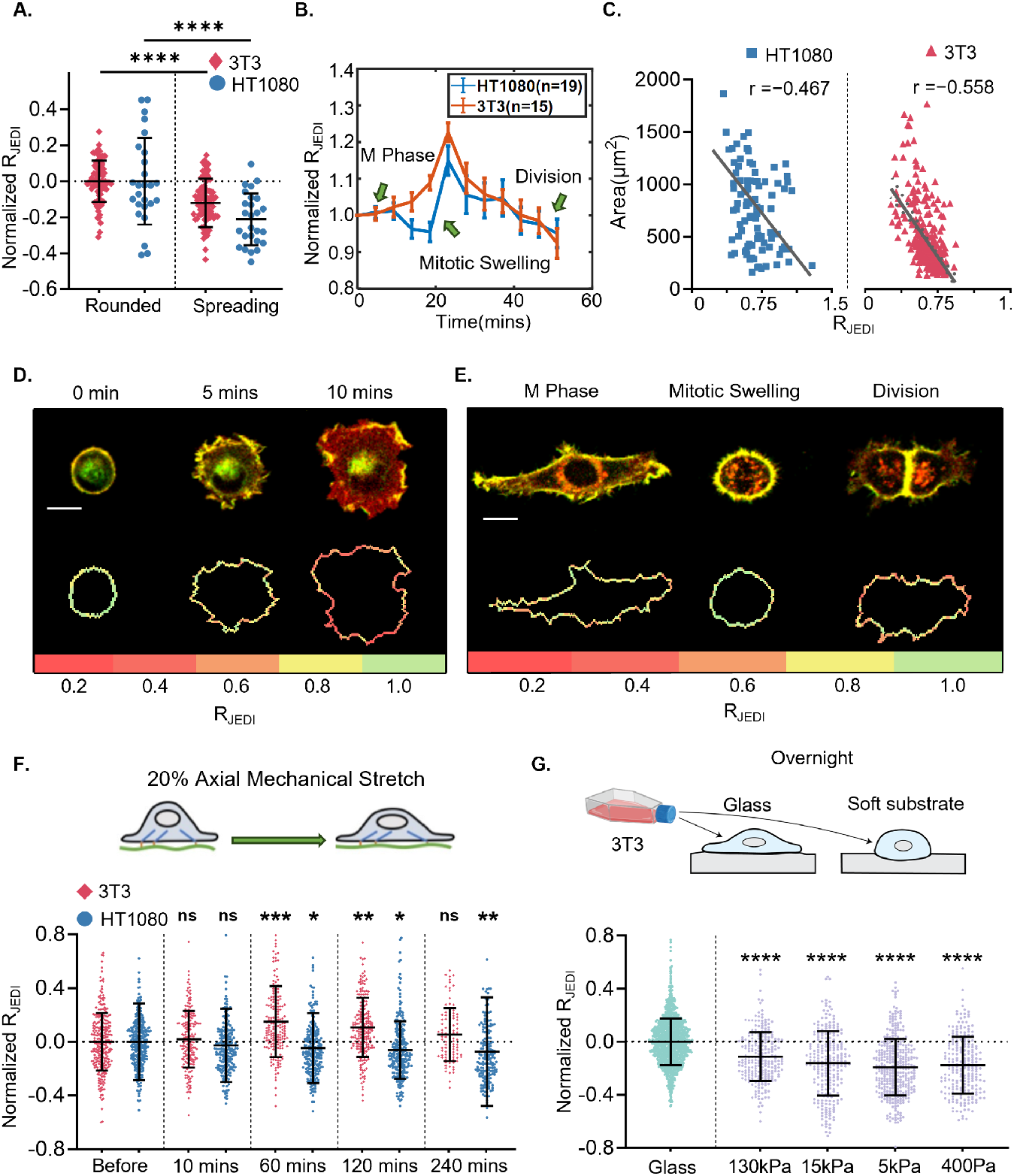
Mechanical stimuli and cytoskeletal dynamics influence cell membrane potential. **(A)** Normalized *R*_JEDI_ in 3T3 and HT1080 before and after cell spreading. *n*_3T3_ = 82. *n*_HT1080_ = 26. **(B)** Time course of normalized *R*_JEDI_ in 3T3 and HT1080 cells during progression from M phase to division. Error bars represent SEM. **(C)** Scatter plots of *R*_JEDI_ versus cell area for HT1080 and 3T3 cells. Pearson correlation coefficients (*r*) are shown. *n*_3T3_ = 100. *n*_HT1080_ = 100. **(D)** Representative confocal images of 3T3 cells expressing JEDI-2P-cyOFP1 during cell spreading, showing *R*_JEDI_ at the cell boundary from rounded (0 min) to a fully spread (10 min) morphology. **(E)** Representative time-lapse confocal images of 3T3 cells during M phase, mitotic swelling, and division into daughter cells, with *R*_JEDI_ mapped at the cell boundary. **(F)** Normalized *R*_JEDI_ in 3T3 and HT1080 cells before and after 20% axial stretch. *n*_3T3_ = 293, 188, 193, 205, 96. *n*_HT1080_ = 281, 204, 237, 238, 185. **(G)** Normalized *R*_JEDI_ in 3T3 cells cultured on glass or soft PDMS substrates overnight. *n*_3T3_ = 1424, 245, 245, 367, 226. **(A, F, G)** Error bars indicate the standard deviation. (A) MannWhitney tests for two-condition comparisons. (F, G) Kruskal-Wallis tests with Dunn’s multiple-comparisons versus pre-stretch (F) or glass (G) condition. **(D, E)** Scale bar: 20 *μ*m.

**Figure S8.**
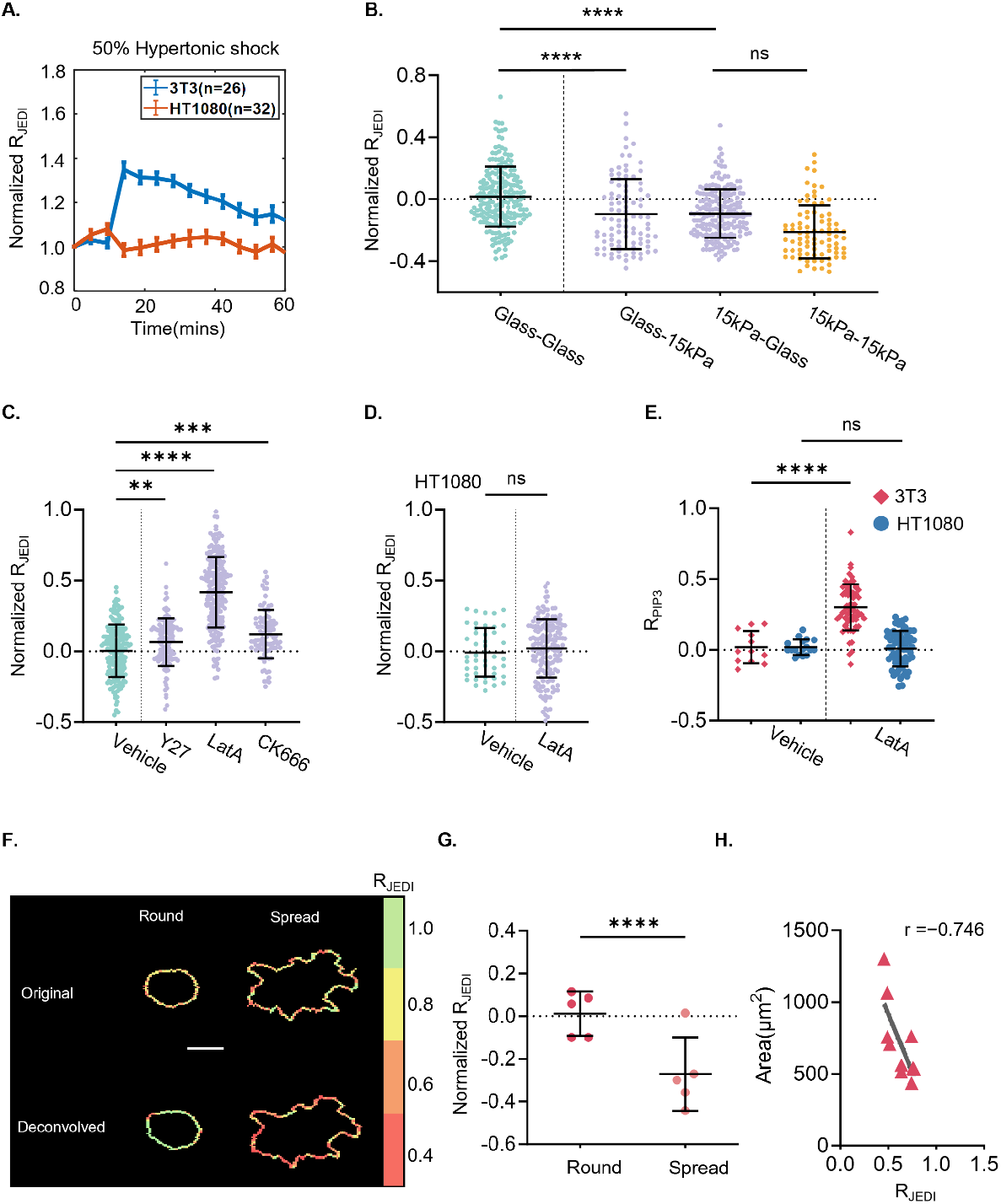
Mechanical stress, substrate stiffness, actin cytoskeleton modulate membrane potential and deconvolved JEDI distribution in round and spread cells. **(A)** Normalized *R*_JEDI_ curve in NIH 3T3 and HT1080 cells under 50% hypertonic shock, showing dynamic membrane potential changes in response to osmotic stress. **(B)** Normalized *R*_JEDI_ changes in NIH 3T3 cells to test the memory effect of soft substrate-induced depolarization. Cells were cultured on glass or 15 kPa soft substrates for five days, then reseeded onto glass or soft substrates, showing that cells originally on soft substrates remained depolarized even when transferred back to glass. *n*_3T3_ = 205, 92, 193, 83. **(C)** Normalized *R*_JEDI_ changes in NIH 3T3 cells after actin cytoskeleton modulation. Treatment with Y27632 (10 *μ*M, a ROCK inhibitor), Latrunculin A (LatA, 2 *μ*M, an actin depolymerization drug), and CK666 (10 *μ*M, an Arp2/3 inhibitor). *n*_3T3_ = 200, 121, 210, 94. **D)** Normalized *R*_JEDI_ changes in HT1080 cells after actin depolymerization using LatA. *n*_HT1080_ = 45, 165. *R*_PIP3_ changes in NIH 3T3 and HT1080 cells after actin depolymerization using LatA. *n*_3T3_ = 12, 65. *n*_HT1080_ = 17, 66. **(F)** Representative confocal images of NIH 3T3 cells expressing JEDI-2P-cyOFP1 during spreading before and after deconvolution, confirming consistency in voltage changes. **(G)** Normalized *R*_JEDI_ in NIH 3T3 cells during spreading after deconvolution, showing the same trend as before deconvolution (Fig. S7A). *n*_3T3_ = 5. **(H)** Scatter plots of *R*_JEDI_ versus cell area in NIH 3T3 cells after deconvolution, showing the same trend as before deconvolution (Fig. S7C). Pearson correlation coefficients (r) are calculated. *n*_3T3_ = 10. **(A)** Error bars represent the SEM. **(B, C, D, E, G)** Error bars indicate the standard deviation. (B, C) Kruskal-Wallis tests followed by Dunn’s multiple-comparisons test were conducted between datasets and the glass-glass condition (B) or the vehicle control (C) of the corresponding cell type. (D, E, G) Mann-Whitney tests were used for two-condition comparisons. **(F)** Scale bar: 20 *μ*m.

## PNP Model of Electrical Potential Across Plasma Membrane

To describe the chemo-electrical field around the cell membrane, in particular the observed spatial gradients in membrane potential, we develop a novel Poisson-Nernst-Planck (PNP) model that includes mobile ions and anionic lipids [1, 3, 2]. The model is developed on a rectangular domain containing a segment of cell membrane and intra/extra-cellular space (Fig. 1). We denote the length of the membrane segment as 2*L* and the thickness as 2*h*. The general PNP equation can be written as:

**Figure 1:**
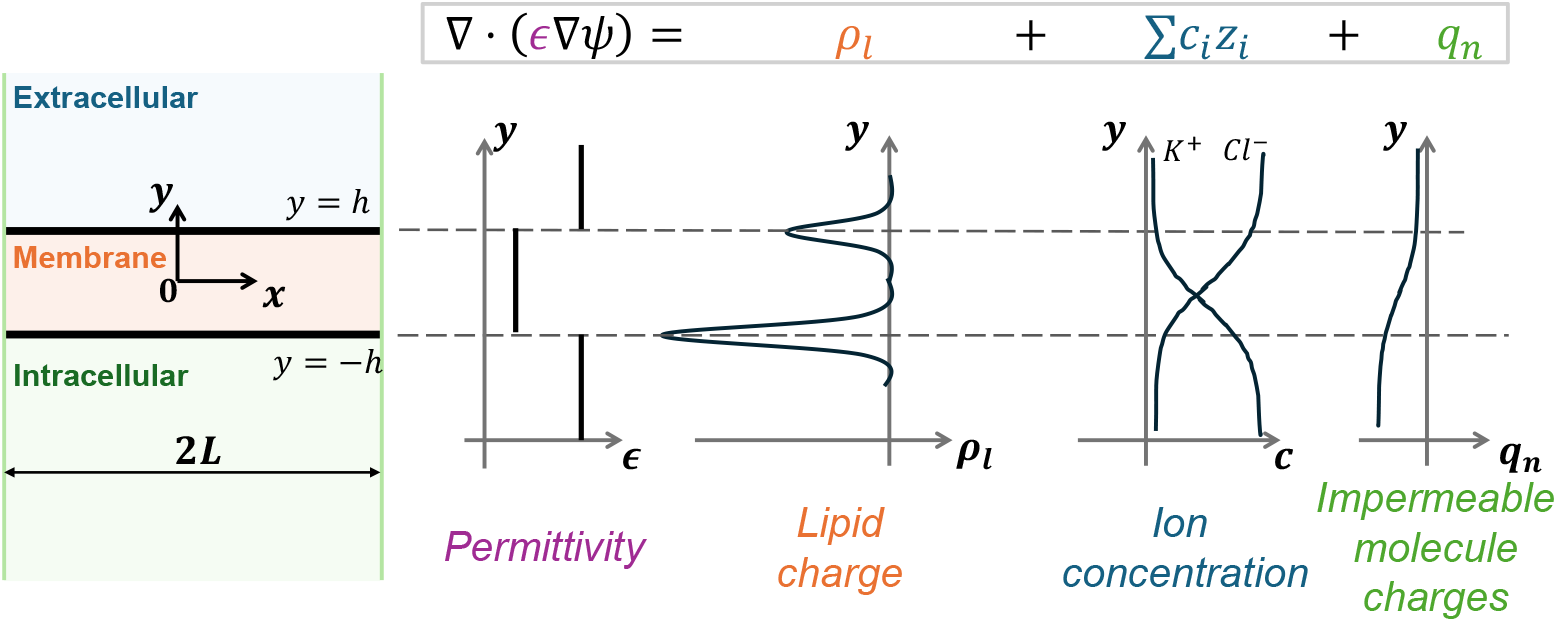
Schematic diagram for the Poisson-Nernst-Planck model. The membrane region (orange) is in between the extracellular and intracellular space. Ions can diffuse in the extracellular and intracellular space, but can only cross the membrane via ion channels/exchangers. Within the membrane, there are also charged lipids: *ρ*_*l*_. The membrane also introduces an oily region with a different electrical permittivity, *ε*. The *y*-dependence of *E* and *ρ*_*l*_ are shown. The model can also include other impermeable charges in the domain.

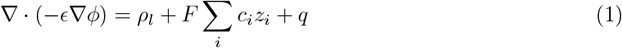

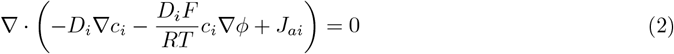

where *φ* is the electrical potential field, *c*_*i*_(*i* = 1, 2, 3) is the concentration of different ions. The valence of different ionic species is denoted as *z*_*i*_. In our model, for simplicity, we consider generic positive and negative ions, modeling K^+^ and Cl^*−*^ (*i* = 1, 2). We also assume that Na^+^ concentration is constant inside and outside the cell, following a step function: *c*_3_ = *c*_3*i*_(*y <*= *−h*), *c*_3_ = *c*_3*o*_(*y > −h*). The active fluxes of these ions are denoted as *J*_*ai*_ (positive along positive direction of the y-axis) and the charge of impermeable anions (e.g., protein, DNA, etc.) is denoted as *q*. Moreover, we also include the lipid charge on the membrane *ρ*_*l*_, which is concentrated at *y* = *−h* (inner leaflet) and *y* = *h* (outer leaflet). Notably, inner leaflet contains more lipid charge than the outer leaflet. The permittivity of different compartments and diffusion coefficients of different ions are demoted as *ε* and *D*_*i*_, respectively. *ε, D*_*i*_ and *J*_*ai*_ are all piece-wise functions with different values in different compartments. *F* is the Faraday constant, *R, T* are gas constant and temperature, respectively. Since we are interested in the steady state, we neglect the time derivatives in Nernst-Planck equation (2). Equations above can be further nondimensionalized as:

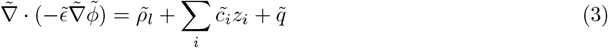

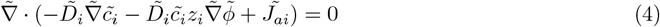

Where the dimensionless parameters are defined as: 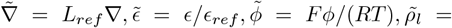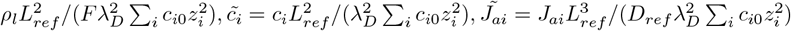. Here *L*_*ref*_, *D*_*ref*_, *ε*_*ref*_ are reference length, diffusion coefficient and permittivity. 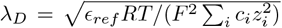 is the Debye length. The Debye length is also a “screening” length where diffusible ions can be influenced by membrane charges. For simplicity, in the following parts, we neglect the tilde in dimensionless parameters.

In order to solve the equations, we also need boundary conditions for the electrical potential and ion concentrations. We here set the external electrical potential as zero in the external bulk environment and assume that it is constant in the intracellular space. This can be written as: *φ*(*y* = ∞) = 0, *∂φ/∂y*(*y* = *−*∞) = 0. Similarly, we assume *i*-th external ion concentration is constant (*c*_*io*_) and intracellular ion concentrations are constant, which is *c*_*i*_(*y* = ∞) = *c*_*io*_, *∂c*_*i*_*/∂y*(*y* = *−*∞) = 0. Periodic boundary conditions are applied along the membrane in the *x*-direction. Values for all parameters are listed in Table 1.

**Table 1:** Glossary of model variables and values in simulation. See text for references.

| Symbol | Description | Value in simulation |
| --- | --- | --- |
| $F$ | Faraday constant (A/mol) | $9.65 \times 10^4$ |
| $R$ | gas constant ($\text{J} \cdot \text{K}^{-1} \cdot \text{mol}^{-1}$) | 8.31 |
| $T$ | temperature (K) | 300 |
| $L_{ref}$ | reference length (nm) | 5 |
| $\epsilon_{ref}$ | reference permittivity (F/m) | $7.04 \times 10^{-10}$ |
| $D_{ref}$ | reference diffusion coefficient ($\text{m}^2/\text{s}$) | $2 \times 10^{-9}$ |
| $c_{1o}$ | extracellular $\text{K}^+$ concentration (mM) | 5 |
| $c_{2o}$ | extracellular $\text{Cl}^-$ concentration (mM) | 145 |
| $c_{3o}$ | extracellular $\text{Na}^+$ concentration (mM) | 140 |
| $c_{3i}$ | intracellular $\text{Na}^+$ concentration (mM) | 15 |
| $\tilde{L}$ | dimensionless length of half of the membrane segment | 5 |
| $\tilde{h}$ | half of dimensionless membrane thickness | 0.5 |
| $\tilde{\epsilon}_i$ | dimensionless permittivity in intracellular space | 1 |
| $\tilde{\epsilon}_m$ | dimensionless permittivity in the membrane | 0.05 |
| $\tilde{\epsilon}_o$ | dimensionless permittivity in extracellular space | 1 |
| $\tilde{q}_i$ | dimensionless charge of nonpenetrating anions in cytoplasm | 20.54 |
| $\tilde{D}_1$ | dimensionless diffusion coefficient of $\text{K}^+$ in intra/extra-cellular space | 1 |
| $\tilde{D}_{1m}$ | dimensionless diffusion coefficient of $\text{K}^+$ in the membrane | 0.5 |
| $\tilde{D}_2$ | dimensionless diffusion coefficient of $\text{Cl}^-$ in intra/extra-cellular space | 1 |
| $\tilde{D}_{2m}$ | dimensionless diffusion coefficient of $\text{Cl}^-$ in the membrane | 0.25 |

**Table 2:**
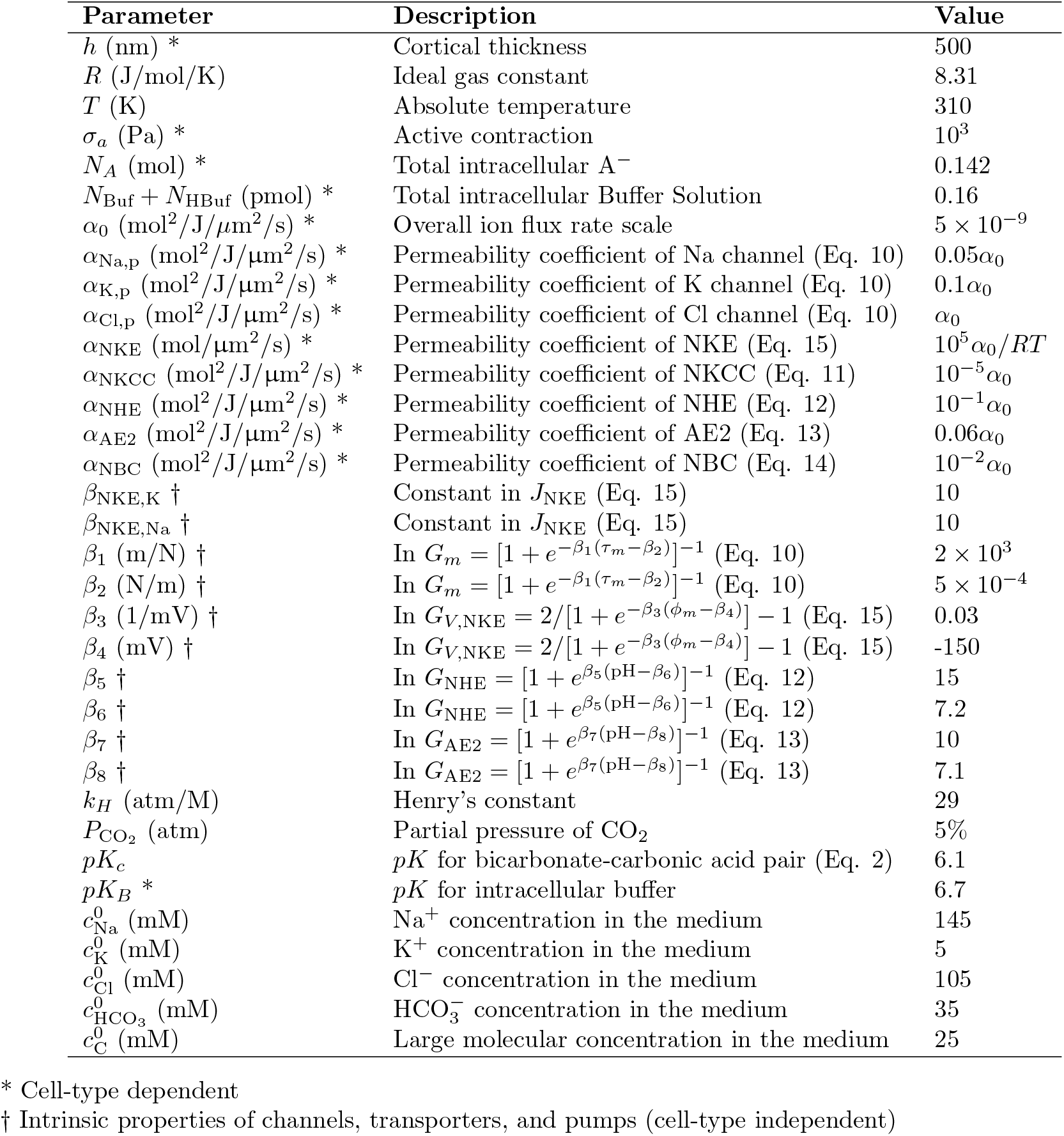
Model parameters.

A typical profile of electrical potential is shown in Fig. 2H in the main text. Notice that there are two different types of membrane potential: LOCAL membrane potential and DISTAL membrane potential. The first type of membrane potential refers to the potential difference between the inner leaflet and outer leaflet while the second type refers to the difference between the intracellular bulk environment and extracellular bulk environment. Both lipid charges and ionic environment will influence the membrane potential profile. We then explore how these two factors influence the electrical potential distribution.

We first examine the case with uniform lipid charge and active ion fluxes. We obtain different potential profiles with different values of *J*_*ai*_ and inner leaflet lipid charge *ρ*_*l*_ (Fig. 2H). Interestingly, lipid charge only contributes to the local membrane potential profile. In contrast, active ion fluxes contribute to both local and distal potential profile. We further introduce spatial variation in lipid charge and active ion fluxes in the *x*-direction. and find that variations in the membrane potential in the *x*-direction is related to the lipid charge spatial variation rather than ion fluxes. These findings quantitatively explain the experimental observation that PIP3 and PS distributions coincide with the membrane potential distribution.

### Relationship Between Membrane Voltage and Ion Exchanger Flux

Our experiments show that ion exchangers such as NHE1 can dramatically influence the membrane potential. We present a cell ion homeostasis model that demonstrates how this phenomenon occurs [4]. The model incorporates ion fluxes, intracellular osmolarity, and pH on the time scale of minutes. We consider the following intracellular species: 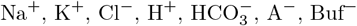, and HBuf, where A^*−*^ represents charged organic molecules or proteins that are not permeable across the cell membrane. The total molar content of intracellular proteins, *N*_*A*_, is prescribed. The organic molecules or proteins have various charges, and the average charge is negative [5]. We thus lump all the proteins and assume an effective average valence of *−*1. Within the cell, we include an unprotonated buffer (Buf^*−*^) and protonated buffer (HBuf) species, both of which are considered as non-permeable across the cell membrane [6], and the total molar content, *N*_Buf_ + *N*_HBuf_, is fixed. The extracellular ion concentrations, 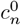, are given based on generic cell culture medium composition or extracellular fluid environment in vivo. 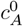 is adjusted based on the concentration of other charged ion species so that the electro-neutrality condition of the medium is satisfied. Here *c* represents concentrations, and the superscript ‘0’ indicates quantities associated with the extracellular environment. We can also include large, non-permeable molecules (G) in the extracellular space to adjust the baseline osmolarity of the medium, in a similar manner as modulating the extracellular hydrostatic pressure, which is determined up to a constant. By default, we let this molecule to be charge-free, but it can assume any valence to ensure the electroneutrality of the extracellular space when other extracellular species are specified. Since the molecule is non-permeable and the electroneutrality condition in the extracellular domain is maintained, it will not affect the membrane potential nor voltage-dependent ion channels, transporters, and pumps.

Considering a generic spherical cell experiencing volume regulation, the rate change of the cell radius, *r*, is modulated by osmotic and hydrostatic pressure gradients [7]

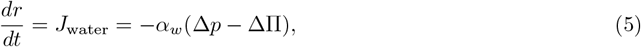

where *J*_water_ is water flux (length per unit time) across the cell membrane, defined positive inward. *α*_*w*_ is the combined permeability coefficient of water from the lipid membrane and aquaporins (AQPs). Since the water permeability through lipids is several orders of magnitude smaller than that of AQPs,*α*_*w*_ is dominated by the contribution from AQPs. 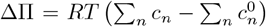 is the osmotic pressure difference across the cell membrane. The extracellular and intracellular osmolytes in the model are, respectively, 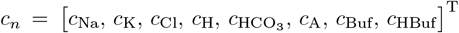 and 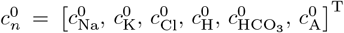 The hydrostatic pressure difference across the cell membrane, Δ*p*, is related to the combined effective cortical and membrane stress, *σ*, by the Laplace law Δ*p* = 2*hσ/r*, where *h* is the combined effective thickness of the cell cortex and membrane. The stress can be obtained from the constitutive relation of the actomyosin cortex and membrane[8] and is composed of two parts, *σ* = *σ*_*p*_ + *σ*_*a*_, where *σ*_*p*_ is the passive mechanical stress of the actin network and membrane, which includes contributions from F-actin crosslinkers, filament mechanics, and lipid bilayer. *σ*_*a*_ is the active myosin contraction from the cortex. The passive stress is typically small compared to the active stress, and in this model, we only consider the active stress such that *σ* ≃ *σ*_*a*_. Hereafter, we will use cortical stress to refer to the combined stress from the cortical layer and the cell membrane. Similarly, we will use cortical tension and cortical thickness to refer to the combined effects from both the cortical layer and the lipid bilayer.

The chemical equilibrium equation for the bicarbonate-carbonic acid pair is 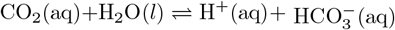, where [CO_2_]_aq_ is related to the partial pressure of 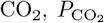, by the Henry constant *k*_*H*_, i.e., 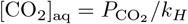. The reaction equilibrium constant for the bicarbonatecarbonic acid pair is 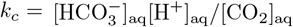. Defining pH_0_ = *−* log_10_[H^+^]_aq,0_ as the extracelluar pH and p*K*_*c*_ = *−* log_10_ *k*_*c*_, we then have 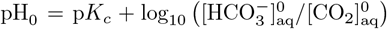, where 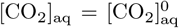 since CO_2_ can move freely across the cell membrane [9]. For the intracellular domain, we have

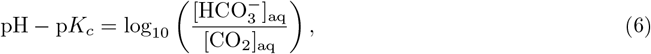

where pH = *−* log_10_[H^+^]_aq_ is the intracelluar pH.

The cell contains various buffer solutions. In general, the chemical reaction for the intracellular buffer solution can be written as H_2_Buf(aq) ⇋ *ℓ*H^+^(aq) + Buf^*ℓ−*^(aq), where *ℓ* = 1, 2, 3,… for different buffer species and *Z*_Buf_ = *− ℓ* is the valence of the unprotonated buffer. The reaction equilibrium constant is similarly defined as 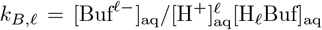. With p*K*_*B, ℓ*_ = *−* log_10_ *k*_*B, ℓ*_,, we have *ℓ*pH *−* p*K*_*B, ℓ*_ = log_10_ ([Buf^*ℓ−*^]_aq_*/*[H_*ℓ*_ Buf]_aq_). Without loss of generality, we lump all buffer solutions into one and use *ℓ*, or equivalently *Z*_Buf_, as a model parameter. By default, we let *ℓ* = 1 and simplify the notation of p*K*_*B*,_, Buf^*ℓ−*^, and H_ℓ_ Buf as p*K*_*B*_, HBuf^*−*^, and HBuf, respectively. Using a different *ℓ* does not change our model conclusion [4].

The conservation equation for the non-reactive ion species is

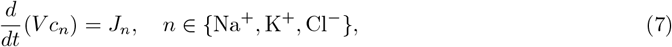

where *J*_*n*_ is the total ion flux (mole per unit time) across the membrane for each species and is determined by the boundary conditions of ion fluxes through the membrane channels, transporters, and pumps; *V* = 4*πr*^3^*/*3 is the cell volume. The conservation equation for the intracellular reactive species is

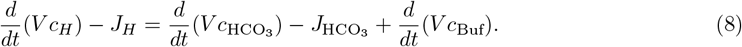

The total flux for each ionic species is the sum of the fluxes through the relevant channels, transports, and pumps:

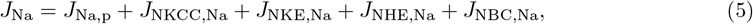

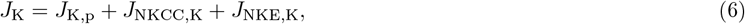

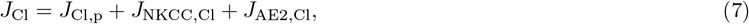

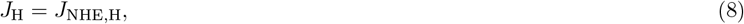

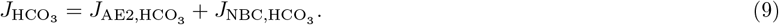

The passive ion fluxes, *J*_*n,p*_, are proportional to the electrochemical potential difference of ions across the membrane [10] and are typically mechanosensitive,

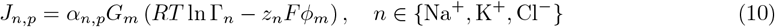

where *φ*_*m*_ is the membrane potential sensed by membrane ion channels; *z*_*n*_ is the valence of each ionic species; 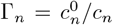 is the ratio of extrato intra-cellular ion concentrations; *α*_*n,p*_ is the permeability coefficient of each species, which depends on the channel property and the density of the channels in the membrane; *G*_*m*_ ∈ (0, 1) is a mechanosensitive gating function that generally follows a Boltzmann distribution, i.e., 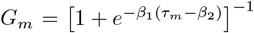, where *β* and *β* are two constants and *τ* is the cortical tension given by *τ*_*m*_ = *σh*.

Since NKCC is mainly a passive transport, the flux through it can be expressed as [11]

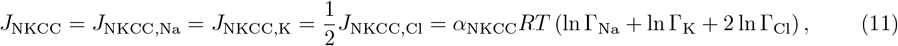

where *α*_NKCC_ is the permeability coefficient independent of the cortical tension. Given the dependence of NHE on pH, the flux of NHE is expressed as

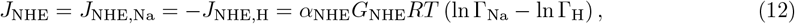

where *α*_NHE_ is the permeability coefficient which does not significantly depend on cortical tension and we assume it as constant. 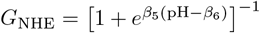 is a pH-gated function indicating the dependence of the NHE activity on pH. Similarly, the flux through AE2 takes the form

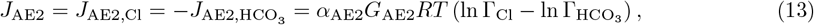

where *α*_AE2_ is the permeability coefficient of AE2 and is assumed to be independent of the cortical tension. 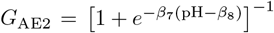 is a pH-gated function indicating the dependence of the AE2 activity on pH. The fluxes through NBC depends on the combined electrochemical potential of the two species, i.e.,

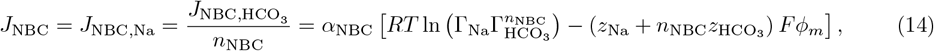

where *α*_NBC_ is the permeability coefficient of NBC and *n*_NBC_ = 2 or 3 indicates the stoichiometry of NBC. We will take *n*_NBC_ = 2 in this work. Since the NKE is voltage-dependent and its flux saturates at high concentration limits, we model the flux of Na^+^ and K^+^ through the Na^+^*/*K^+^ pump as

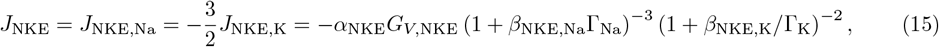

where *α*_NKE_ is the permeability coefficient of the pump depending on the density of the pump as well as the concentration of ATP. *β*_NKE,Na_ and *β*_NKE,K_ are constants that scale Γ_Na_ and Γ_K_, respectively. The exponents 3 and 2 are the Hill’s coefficients of Na^+^ and K^+^, respectively. Equation 15 ensures that the flux is zero when either 1*/*Γ_Na_ or Γ_K_ approaches zero; the flux saturates if 1*/*Γ_Na_ and Γ_K_ approaches infinity. *G*_*V*,NKE_ captures the voltage-dependence of the pump activity [12], 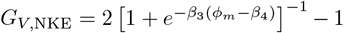, where *β*_3_ and *β*_4_ are constants. The electro-neutral condition should be satisfied for both the intracellular and the extracellular spaces. The condition for the intracellular space is maintained by enforcing ∑*z*_*n*_*c*_*n*_ = 0. The full set of equations for the system is as follows,

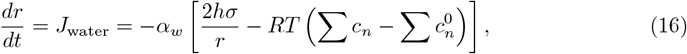

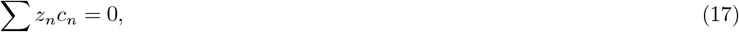

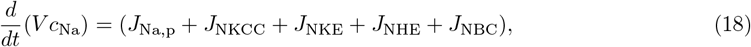

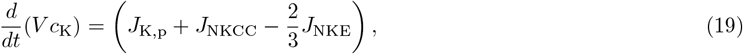

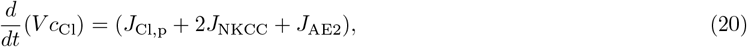

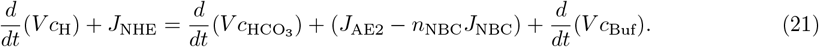

The given quantities (known model inputs or parameters) are: 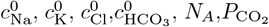, and *N*_HBuf_ + *N*_Buf_. The six equations (Eqs. 16 to 21) are used to solve for the six unknowns: *r, φ*_*m*_,*c*_Na_, *c*_K_, *c*_Cl_, and pH. The rest of the quantities are derived either from the given quantities or from the unknowns.

For steady-state response, Eqs. 16 to 21 can be directly implemented to solve for the six unknowns **x** = (*r, φ*_*m*_, *c*_Na_, *c*_K_, *c*_Cl_, pH)^T^ by letting *d/dt* = 0. We will use a steady-state model to study the effect of membrane ion channel inhibitors on cell homeostatic. The model parameters are provided in Tab. 2.

The model predicts that inhibition of NHE, AE2, or NKCC leads to membrane hyperpolarization, while inhibition of NKE results in depolarization (Fig. 2), consistent with experimental observations. In this context, the membrane potential refers to the voltage difference that drives ionic fluxes necessary to maintain intracellular electroneutrality. Importantly, this potential represents the local voltage sensed by ion channels, rather than the bulk cytoplasmic potential. Although the model does not explicitly account for the lipid composition that contributes to the local membrane environment, it does predict the membrane potential required to support the predicted intracellular ion concentrations and membrane ion fluxes (Fig. 2).

**Figure 2:**
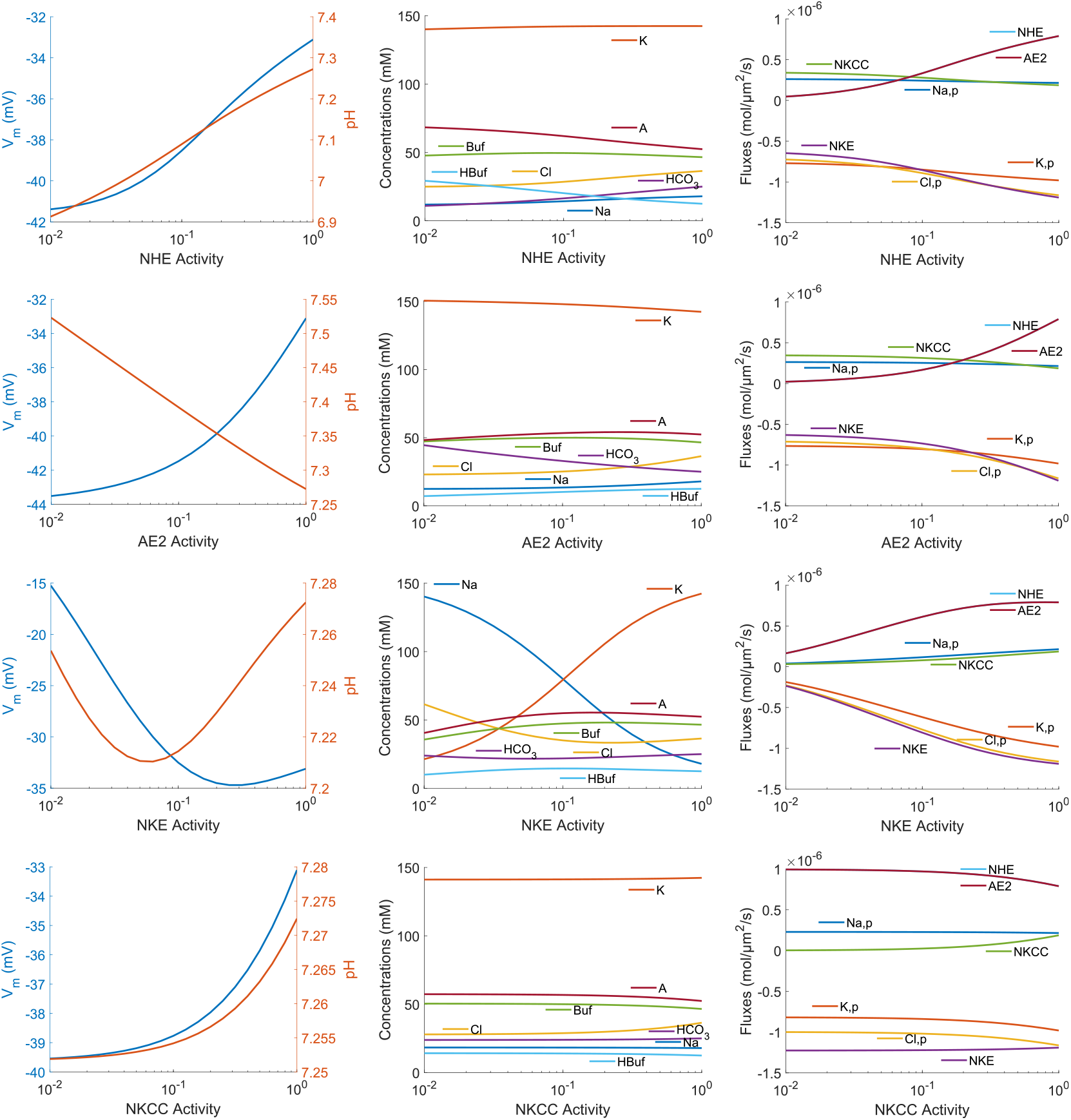
Model prediction on the membrane potential, intracellular pH, ion concentrations, and ionic fluxes upon various ion transporter inhibition.

**Figure 3:**
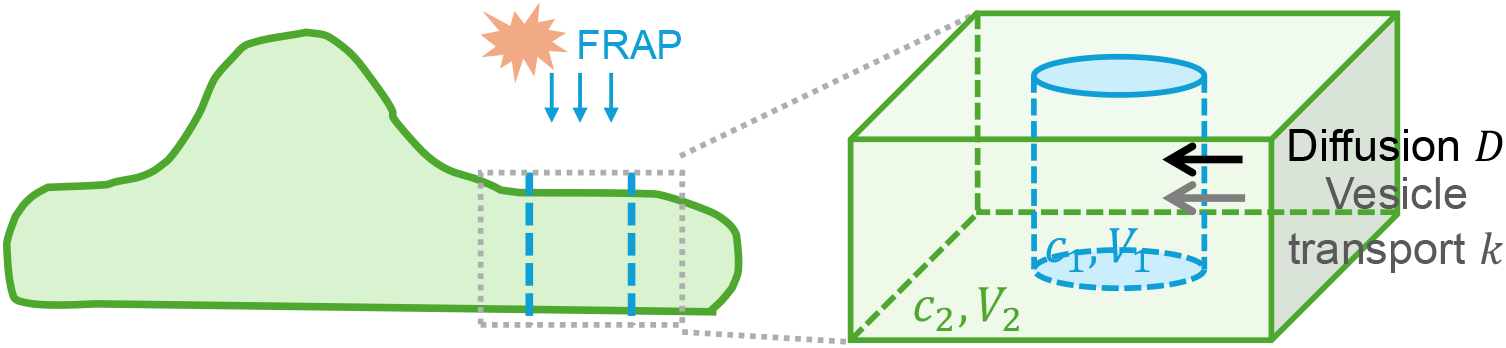
Schematic diagram for the FRAP recovery model. In the experiment, a subvolume of the cytoplasm is bleached by laser. The intensity of Lact-C2 is followed over time. The intensity in the FRAPed region recovers from fluorophore diffusion as well as vesicle transport into the region.

### Modeling of Anionic Lipid Transport and FRAP Intensity Recovery

To interpret the FRAP experiment results of the phosphatidylserine (PS) indicator Lact-C2 (Fig. 4 in the main text), we developed a dynamical model of photobleaching recovery that incorporates both diffusion and vesicle-mediated transport of Lact-C2 following photobleaching (Fig. S3).

We consider two cytoplasmic regions: the *FRAP region* and the *external region*, with volumes denoted by *V*_1_ and *V*_2_, respectively. The concentrations of Lact-C2 in these two regions are represented by *c*_1_ and *c*_2_. Before photobleaching, Lact-C2 is assumed to be uniformly distributed such that

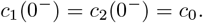

Immediately after bleaching, the concentrations become *c*_1_(0^+^) and *c*_2_(0^+^), where *c*_1_(0^+^) *< c*_0_ and *c*_2_(0^+^) = *c*_0_.

After photobleaching, the recovery of fluorescence in the FRAP region occurs through both diffusion and active vesicle transport from the external region, characterized by the diffusion coefficient *D* and the active transport rate constant *k*. The temporal evolution of the concentrations is described by:

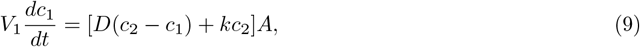

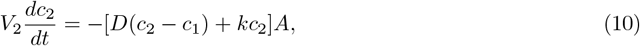

where *A* denotes the contact surface area between the FRAP region and the external environment. For the active transport term, we model the net import of Lact-C2 into the FRAP region, which is proportional to the external concentration *c*_2_ with proportionality constant *k*.

At steady state, the concentrations are denoted by 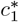 and 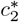, and the FRAP recovery ratio is given by:

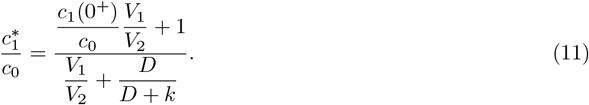

This expression indicates that the steady-state recovery ratio depends on three key factors: (1) the initial bleaching depth, quantified by the ratio *c*_1_(0^+^)*/c*_0_; (2) the volume fraction of the FRAP region, *V*_1_*/V*_2_; and (3) the active import rate constant *k*. The initial bleaching depth *c*_1_(0^+^)*/c*_0_ and the FRAP volume fraction *V*_1_*/V*_2_ are both less than 1. *V*_1_*/V*_2_ 1. Therefore, the leading order estimate for the steady state recovery ratio is

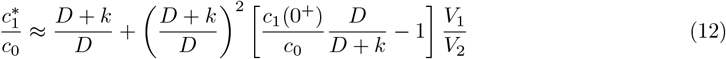

which implies that higher steady state recovery ratio is dominated by larger import flux *k* into the FRAP region. Therefore, our data and the model suggest that the vesicle transport rate *k* is larger in the lateral region than the protrusion region, indicating higher levels of transport activity.

## Notes

### Competing Interest Statement

The authors have declared no competing interest.

